# Tumor Niche Network-Defined Subtypes Predict Immunotherapy Response of Esophageal Squamous Cell Cancer

**DOI:** 10.1101/2023.02.15.528539

**Authors:** Kyung-Pil Ko, Shengzhe Zhang, Yuanjian Huang, Bongjun Kim, Gengyi Zou, Sohee Jun, Jie Zhang, Cecilia Martin, Karen J. Dunbar, Gizem Efe, Anil K. Rustgi, Haiyang Zhang, Hiroshi Nakagawa, Jae-Il Park

## Abstract

Despite the promising outcomes of immune checkpoint blockade (ICB), resistance to ICB presents a new challenge. Therefore, selecting patients for specific ICB applications is crucial for maximizing therapeutic efficacy. Herein we curated 69 human esophageal squamous cell cancer (ESCC) patients’ tumor microenvironment (TME) single-cell transcriptomic datasets to subtype ESCC. Integrative analyses of the cellular network transcriptional signatures of T cells, myeloid cells, and fibroblasts define distinct ESCC subtypes characterized by T cell exhaustion, Interferon (IFN) a/b signaling, TIGIT enrichment, and specific marker genes. Furthermore, this approach classifies ESCC patients into ICB responders and non-responders, as validated by liquid biopsy single-cell transcriptomics. Our study stratifies ESCC patients based on TME transcriptional network, providing novel insights into tumor niche remodeling and predicting ICB responses in ESCC patients.

## Introduction

Esophageal cancer is the seventh most prevalent cancer, and ESCC accounts for more than 80% of esophageal cancer cases worldwide.^1, 2^ The leading cause of cancer death of ESCC is the sixth high of all types of cancer as the 5-year survival rate is as low as 10-25 %.^3^ Despite its high incidence, the treatment option for ESCC is limited compared to the other major types of cancer. Among the multidisciplinary treatment, including surgery, neoadjuvant therapy, and chemoradiotherapy, therapeutic option for ESCC largely relies on cytotoxic reagent-based chemotherapy. However, the outcome is unfavorable.^4, 5^

To overcome the limited efficacy of ESCC treatment, immunotherapy using immune checkpoint inhibitors (ICI) ^6^ has recently been tested in clinical trials, which resulted in survival benefits for advanced or metastatic ESCC patients.^7-9^ However, approximately 34% and 25% of ESCC patients discontinued ICI treatment because of disease progression^9^ and severe adverse effects,^10, 11^ respectively. In recent clinical trials, the ICI response rate of ESCC patients was only 17% to 28%.^10-12^. Although the clinical trials using ICIs are mainly applied to patients diagnosed with advanced or metastatic ESCC, pathologic criteria used for selecting patients for ICIs remain to be clarified.^5^ Despite the modern pathological criteria, such as PD-L1 expression in tumor cells, stratifying ESCC patients for specific ICIs becomes crucial in improving the effectiveness of immunotherapy.

Tumor microenvironment (TME), a cellular niche surrounding tumor cells, includes immune cells, fibroblasts, and endothelial cells. ^13^ Accumulating evidence suggests that TME plays a crucial role in tumor progression, metastasis, therapy resistance, and immune evasion.^14^ Along with the advent of single-cell transcriptomics, the oncogenic functions of TME in ESCC tumorigenesis have been recently unraveled. Several studies characterized ESCC TME as creating an immunosuppressive environment.^15-18^ In addition to the conventional cancer classification, which mainly relies on the pathologic stages,^19^ transcriptome-based cancer classification has recently been introduced in several cancer types.^20, 21^ Simultaneously, profiling cancer immune systems or cancer-associated fibroblasts (CAFs) identified tumorigenic roles of tumor-infiltrated_ immunocytes and CAFs, which also gained attention.^22, 23^ Nonetheless, comprehensive dissection and characterization of ESCC TME still needed to be achieved. Moreover, how distinct TMEs define immune evasion and ICB response of ESCC remains to be determined.

Herein we analyzed 69 single-cell transcriptomic datasets of ESCC patients’ primary tumor samples and characterized whole TME. Intriguingly, comprehensive analyses of TME identified the distinct networks among T cells, myeloid cells, and fibroblasts, which define specific subtypes and immunosuppression of ESCC.

## Results

### Tumor immune environment transcriptome-based classification of ESCC patients

To elucidate the tumor microenvironment (TME)-based patient characterization, we analyzed single-cell RNA-sequencing (scRNA-seq) datasets of ESCC primary tumors from 69 patients (Fig. 1). All datasets were integrated using the Harmony algorithm^24^ and processed to analyze only non-epithelial cells (*EPCAM* negative) comprising ESCC TME (Fig. 2A and 2B). Unsupervised transcriptomic clustering revealed several immune cell types, fibroblasts, and mast cells (Fig. 2C and Supplementary Fig. S1A). Since T cells play a pivotal role in eliciting an immunogenic response to tumor cells, we first analyzed T cell clusters.^25, 26^ T cell clusters were isolated and processed into the subgroups (Fig. 2D). The unsupervised clustering using principal component analysis (PCA) and Pearson’s correlation categorized T cells of 69 datasets into four groups (T1, T2, T3, and T4) (Fig. 2E). On the Uniform Manifold Approximation and Projection (UMAP), T cells of T1 and T3 groups were closely located together. In contrast, the T4 group was slightly distinct from the T1 and T3 groups (Fig. 2F). Notably, the T2 group was the most distantly located on the UMAP, indicating the minor similarity of T2 transcriptome compared to that of T1, T3, and T4 groups (Fig. 2F). To define subsets of T cells, we annotated T cells based on the marker genes expression (Fig. 2G and Supplementary Fig. S1B and S1C). In a detailed cell subset analysis, T cells of the T2 group showed the most abundance in CD8 T cells, while the other subsets (T1, T3, and T4) rarely exhibited CD8 T cells (Fig. 2G). Besides, T cells of the T4 group showed the highest proportion of exhausted T cells (T_ex_) compared to the other three groups (Fig. 2G). Since T2 is the distinct subgroup, we further analyzed the T cells of the T2 group. After excluding the CD4 T cell cluster, we found that T2 group T cells can be classified into several memory T cells based on the marker genes of subsets (Supplementary Fig. S1D and S1E).^27, 28^ Interestingly, late memory T cells and effector T cells were observed to be the most frequent cells in the T2 group compared to the other subsets (Supplementary Fig. S1F). These results indicate that the T4 group of patients is mainly characterized by T cell exhaustion, whereas the T2 group is enriched with active CD8 T cells.

**Figure 1.**
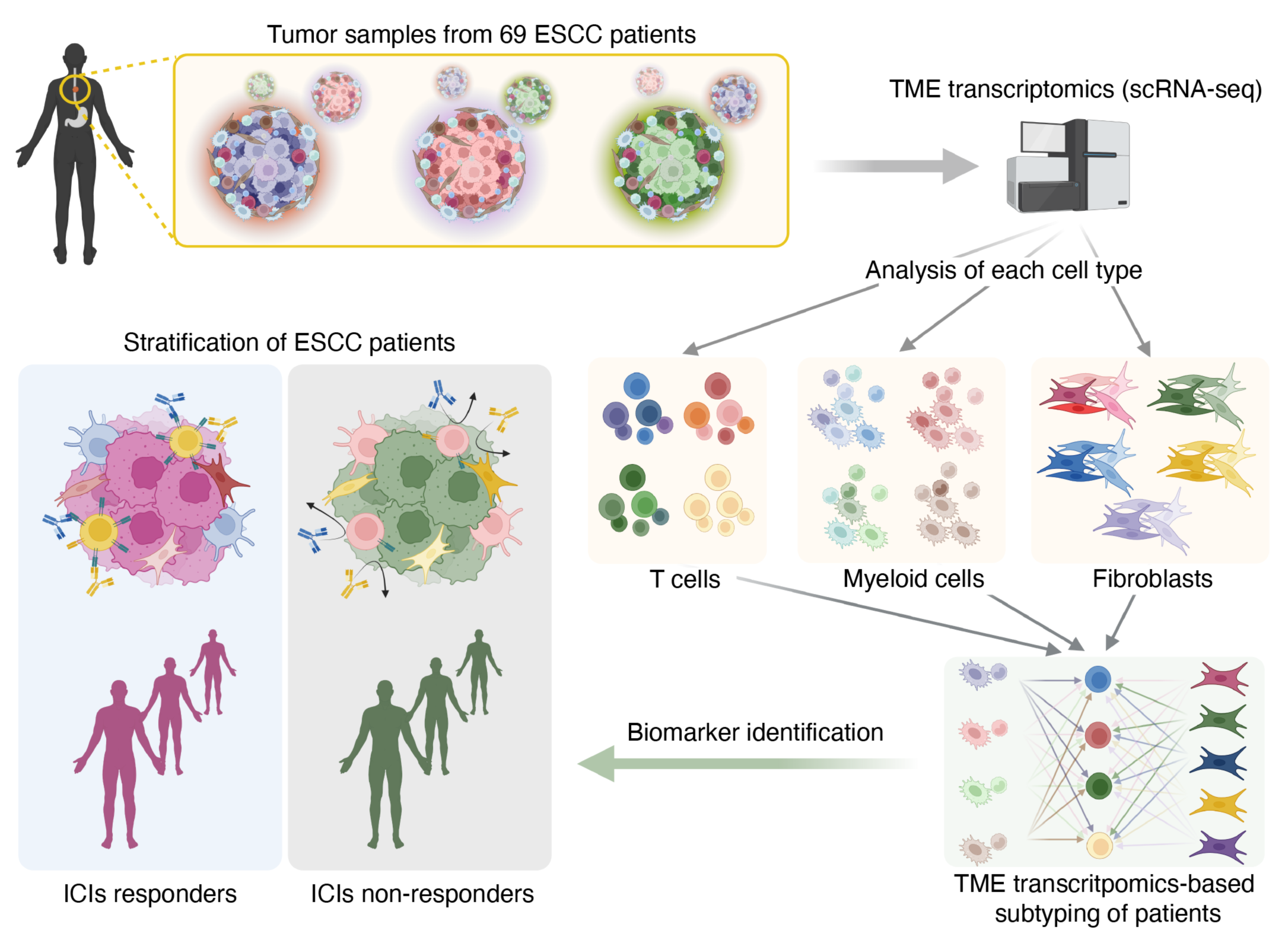
| Schematic workflow for transcriptomic analysis of TME from ESCC patients.

**Figure 2.**
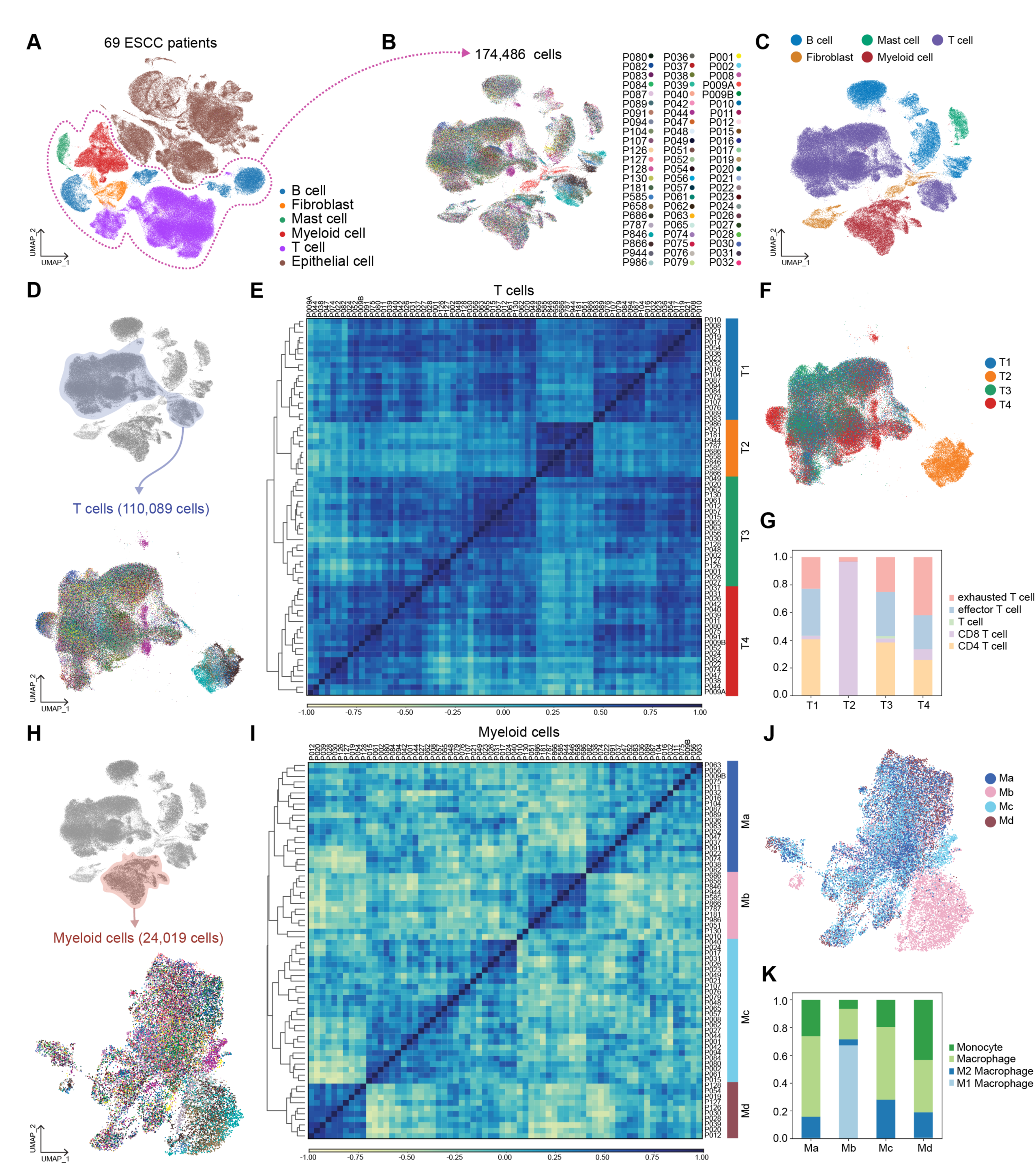
| Immune cells analysis and classification. **A,** Uniform Manifold Approximation and Projection (UMAP) display of whole cells from 69 patients. Single-cell RNA-sequencing (scRNA-seq) results of the cells of TME were integrated and projected **B-C,** Non-epithelial cells were isolated, and UMAP was redrawn with individual patient’s information. (**B**) and five major cell types (**C**). **D,** UMAP display of T cells subgroup with unique patients ID. T cells were isolated from immune cells and clustered again. **E,** T cells were classified into four sub-groups by principal component analysis (PCA) and Pearson correlation. PCA result was clustered by the dendrogram, and Pearson correlation was displayed by color spectrum. **F,** T cells were displayed in UMAP based on the sub-groups defined from PCA and Pearson correlation. **G,** Each sub-groups of T cells were shown with subsets using stacked bar plots. **H,** Myeloid cells of each patient were displayed with UMAP. Myeloid cells were isolated from immune cells and clustered independently. **I,** Myeloid cells were categorized into four sub-groups by PCA and Pearson correlation. PCA results were displayed with a dendrogram, and Pearson correlation was shown by color spectrum. **J,** Myeloid cells were displayed with sub-groups identified from PCA and Pearson correlation. **K,** Each myeloid cell sub-group was displayed with subsets of myeloid cells.

In addition, we comparatively analyzed myeloid cell clusters of 69 datasets (Fig. 2H). Similar to T cell analysis, Myeloid cells transcriptomes were classified into four groups (Ma, Mb, Mc, and Md) based on the principal component analysis and Pearson’s correlation (Fig. 2I). Myeloid cells of Ma and Mc showed close location on the UMAP. In contrast, some Md cells were distinguishable from Ma and Mc. The most distinct myeloid sub-group was Mb on the UMAP (Fig. 2J). Interestingly, based on the clustering with marker genes, Mb-grouped myeloid cells were enriched with M1 macrophages, whereas possessing the least proportion of M2 macrophages compared to the other three groups (Ma, Mc, and Md) (Fig. 2K and Supplementary Fig. S1G). The Ma and Mc groups of myeloid cells were enriched with macrophages. Md groups showed the highest proportion of M2 macrophages among the four groups (Fig. 2K). These results imply that the ESCC patients in the Mb group might have tumor-unfavorable myeloid cells compared to the other groups.

### Single-cell transcriptomes of myeloid and T cells define immunosuppressive ESCC subtypes

We next evaluated which group of T cells exhibits the most immunosuppressive characteristics by the expression of T_ex_ markers. The T4 group expressed the highest level of *LAG3*, *PDCD1*, and *HAVCR2*, whereas the T2 group barely expressed the T_ex_ cell markers (Fig. 3A and Supplementary S2A). To test if the T cell category correlates with myeloid cell classification, we compared the frequency of each patient of the T cell group in the myeloid cell group. Interestingly, patients of T2, the least T_ex_-characterized group, solely belonged to the Mb categories. T4-grouped patients, the enriched T_ex_-characterized group, were mainly distributed to Ma-or Mc-grouped patients (Fig. 3B and 3C). The most frequent patients of T4 were identified as Ma-and Mc-grouped patients. Besides, T1- and T3-grouped patients were primarily directed from Ma- or Mc- grouped patients and Mc- or Md-grouped patients, respectively (Fig. 3B and 3C). Accordingly, we combined the categories of T cells and myeloid cells to make 13 sub-groups (M-T groups) of patients (Fig. 3D). We observed that the Mb-T2 group was separated from other cell clusters in the T cell UMAP. The Ma-T4 or Mc-T4 groups were slightly distinct from the significant population of T cells in the UMAP (Fig. 3E). We also identified that Ma-T4 and Mc-T4 groups exhibited the highest expression of T_ex_ cell markers. Conversely, the Mb-T2 group showed the lowest expression of those markers (Fig. 3F and Supplementary Fig. S2B and S2C). Based on these findings, we analyzed Ma-T4 and Mc-T4 groups of patients with T cells, epithelial cells, and myeloid cells since these groups showed the highest expression of T_ex_ markers in T cells. In the GSEA comparing Ma-T4 or Mc-T4 with the Mb-T2 group of T cells, both Ma-T4 and Mc-T4 groups displayed enrichment of ‘Negative regulation of lymphocyte activation’ and ‘IFN α/β signaling’ (Fig. 3G and 3H and Supplementary Fig. S2D), implying that T cells are enriched with type I IFN signaling in the Ma-T4 and Mc-T4 groups. Since Ma-T4- and Mb-T2-grouped T cells have the most and least T_ex_ features, respectively, we compared these two groups in T cells to identify specific signaling pathways for each group. From the positive signaling in Ma-T4 and negative signaling in Mb-T2 groups, we found that ‘PD-1 signaling’ and ‘IFN α/β signaling’ were shared in T cells of both groups by GSEA analysis (Fig. 3I and 3J). These results suggested that Ma-T4-grouped T cells were relatively enriched with immunosuppressive signaling compared to the Mb-T2- grouped T cells. Additionally, from the GSEA of epithelial and myeloid cells, epithelial cells of Ma-T4 and Mc-T4 groups were observed to show enriched ‘IFN α/β signaling’, consistent with the result from T cell (Supplementary Fig. S2E and S2F). These results suggest that Ma-T4 and Mc-T4 groups are characterized by T cell exhaustion and IFN signaling activation compared to the Mb-T2 group.

**Figure 3.**
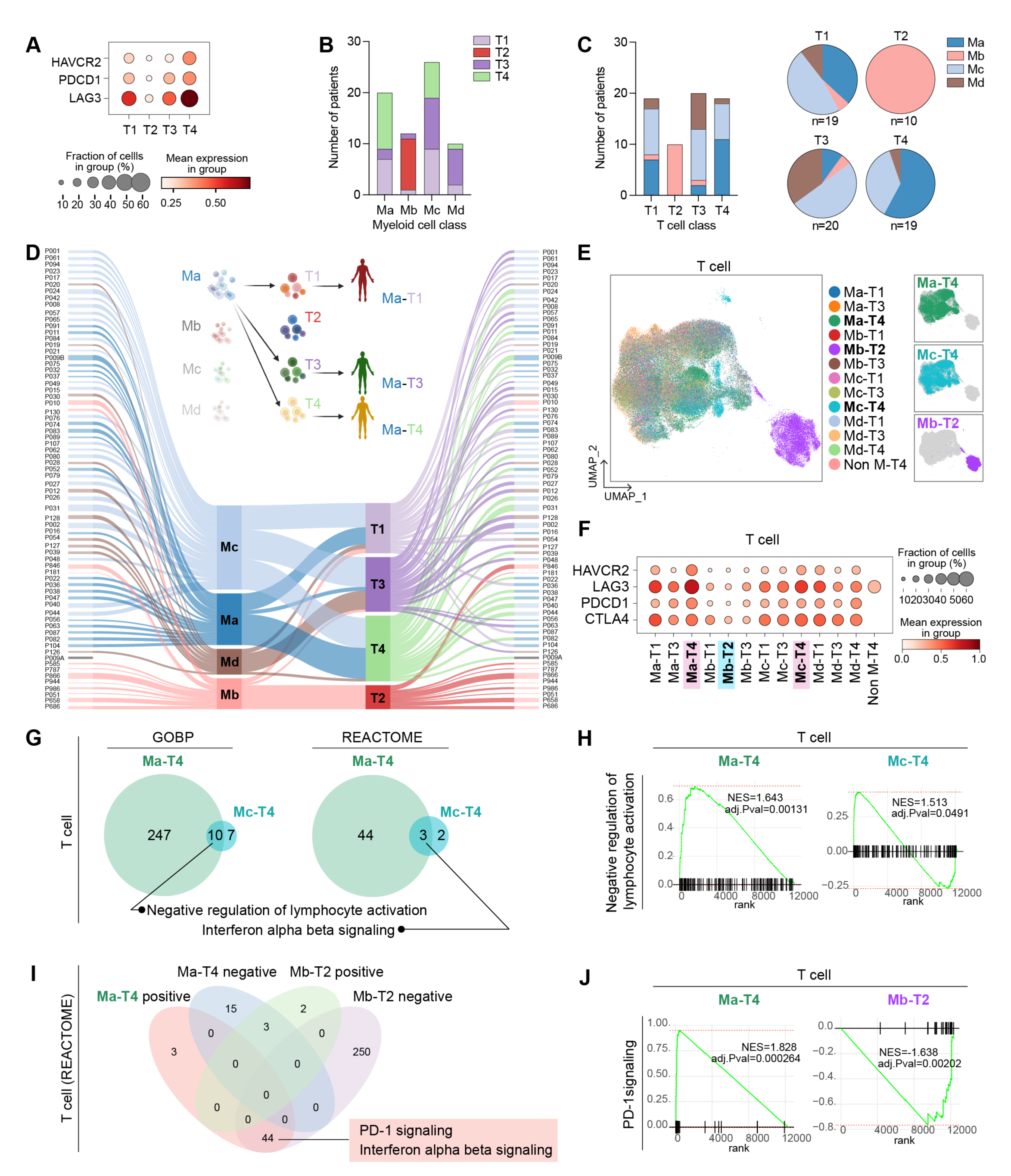
| Comparative analysis of patients by myeloid and T cell classifications. **A,** T_ex_ cell markers expression in each T cell sub-group. **B,** The number of patients of T cell-sub-groups was displayed in each patient’s sub-groups categorized by myeloid cells. **C,** The number of myeloid cell-sub-groups was displayed in each patient’s sub-groups categorized by T cells. The proportion of myeloid-cell-based classified patients in each sub-group of T cells was shown with pie plots. **D,** Individual patients were subjected to each sub-group of myeloid and T cells by Sankey plot. P009A patient was not included in the myeloid cell-based sub-group due to the lack of myeloid cells in the dataset. Each patient was classified into 13 groups (M-T groups) by sub-groups of myeloid cells and T cells and categories. **E,** T_ex_ cell markers expression in T cells in M-T groups of patients. **F,** T cells of each patient from 13 groups were displayed with UMAP. **G-H,** GESA analysis was performed in T cells of M-T groups of patients. The results of GSEA from the Ma-T4 and Mc-T4 groups of patients were compared. GOBP and REACTOME databases were used, and the significant signaling pathways with positive values of NES were compared. Overlapped signaling pathways were displayed with a Venn diagram (**G**) and enrichment plot (**H**). **I**-**J,** GSEA analysis was performed in T cells of Ma-T4 and Mb-T2 patients. significant signaling pathways with both positive and negative valued of NES were compared, and the shared signaling, which has positive values of NES in Ma-T4 and negative values of Mb-T2 were analyzed. The number of shared and exclusive signaling in each group was shown in the Venn diagram (**I**). PD-1 signaling, shared signaling in T cell GSEA analysis of Ma-T4 positive and Mb-T2 negative, was displayed with enrichment plots (**J**).

## Cellular interactome identifies the TIGIT-NECTINE2 pathway as a co-suppressor for immunosuppressive TME

We next performed cell-to-cell interaction analysis using the ‘CellChat’ package that infers cellular interactome based on ligand-receptor expressions.^29^ Comparative analysis of cell-to-cell interactions in Ma-T4 and Mb-T2 identified that TIGIT, NECTIN, and PD-L1 signaling were significantly enriched in the Ma-T4 patient group (Fig. 4A).

**Figure 4.**
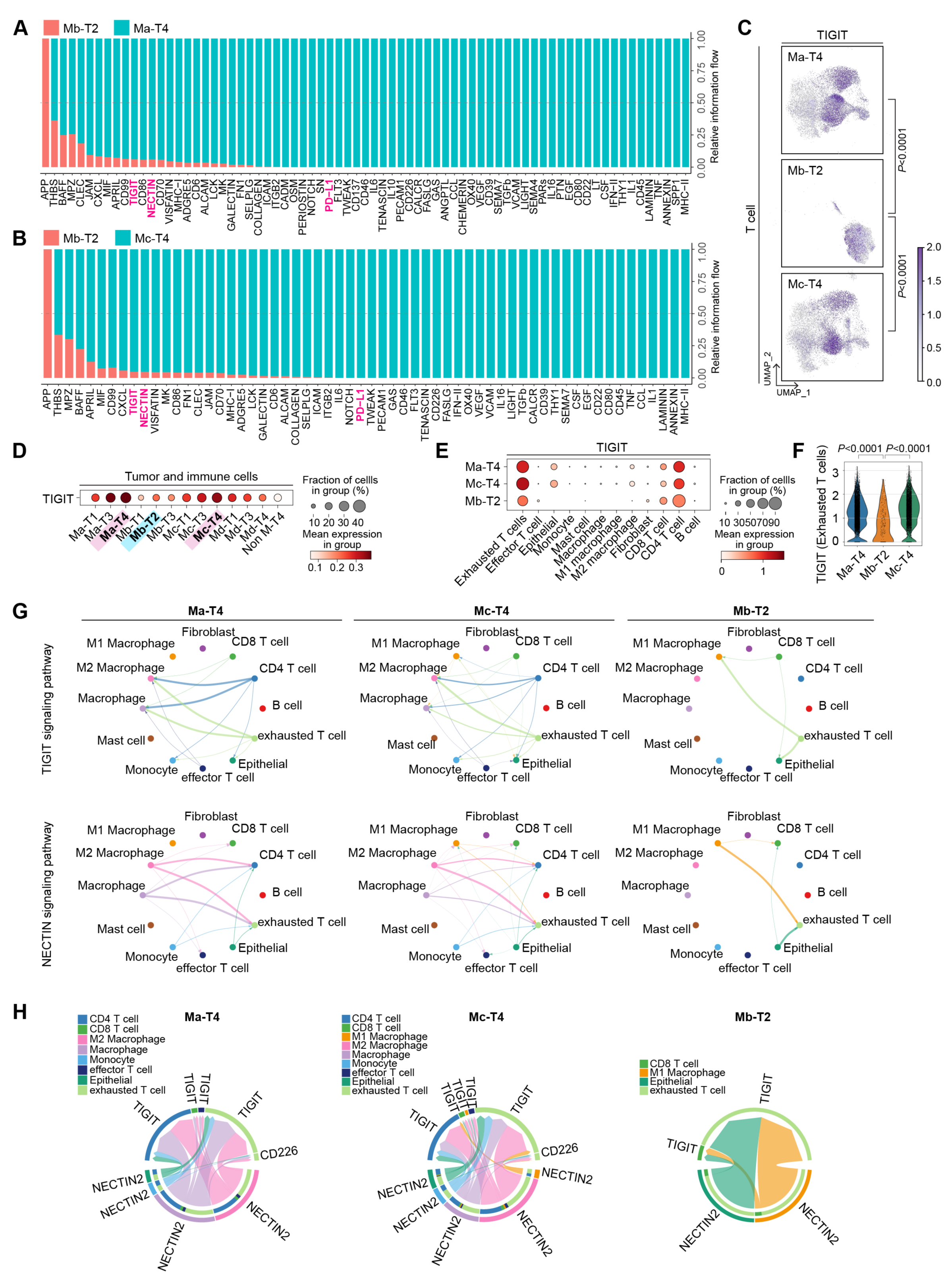
| Cell-to-cell interactions comparison in M-T groups of patients. **A,** Enriched cell-to-cell signaling calculated by CellChat was compared in the Ma-T4 and Mb-T2 group of patients. T cell exhaustion-related signaling pathways were highlighted. **B,** Enriched cell-to-cell signaling calculated by CellChat was compared in the Mc-T4 and Mb-T2 group of patients. T cell exhaustion-related signaling pathways were highlighted. **C,** TIGIT expression in the T cells was displayed with feature plots. T cells of Ma-T4, Mc-T4, and Mb-T2 groups were separated and projected. **D,** TIGIT expression in M-T groups was shown with a dot plot. All the cells, including tumor and immune cells, were compared in each group of patients. **E,** TIGIT expression in each cell type was compared. Ma-T4, Mc-T4, and Mb-T2 groups of patients were displayed. **F,** TIGIT expression in T_ex_ cells was compared in Ma-T4, Mb-T2, and Mc-T4 sub-groups. **G,** Significant interactions within cell types were shown with circle plots. TIGIT and NECTIN signaling pathways were compared in the Ma-T4, Mc-T4, and Mb-T2 groups of patients. **H,** Specific genes related to NECTIN signaling pathways were displayed with chord plots. The source group of cell types was located on the bottom hemispheres, and the receiver group was on the top hemispheres. Ma-T4, Mc-T4, and Mb-T2 groups of patients were compared.

The same results were also observed in the comparison between Mc-T4 and Mb-T2 groups of patients (Fig. 4B). Consistently, TIGIT expression was higher in Ma-T4 and Mc-T4 groups while lower in Mb-T2 group, especially in T cells (Fig. 4C and 4D and Supplementary Fig. S3A). TIGIT was primarily expressed in T_ex_ and CD4 T cells as previously reported (Fig. 4E).^30, 31^ The expression of TIGIT in T_ex_ was markedly higher in Ma-T4 and Mc-T4 groups compared to the Mb-T2 group (Fig. 4F). As NECTIN2 is known to be a ligand for TIGIT and CD226,^32^ TIGIT and NECTIN2 signaling-mediated interactions were significant and abundant in Ma-T4- and Mc-T4-grouped patients compared to Mb-T2 patients with similar interacting patterns (Fig. 4G). Moreover, NECTIN2 was expressed mainly in epithelial cells, which implies possible interaction between epithelial cells and T cells through NECTIN2 and TIGIT (Supplementary Fig. S3C and S3D). However, the expression of NECTIN2, a competitive ligand of TIGIT and CD226, was not significantly higher in Ma-T4 or Mc-T4 group compared to the other groups (Supplementary Fig. S3E). We analyzed the specific genes in cell-to-cell interactions and found that various types of cells, including epithelial and myeloid cells, were predicted to interact with T cells and myeloid cells via NECTIN2 and TIGIT. The interactions between NECTIN2 and TIGIT or CD226 were more abundant in Ma-T4 and Mc-T4 groups compared to Mb-T2 (Fig. 4H). These results suggest that T cell activation inhibitory signaling, i.e., NECTIC and TIGIT, could be therapeutic targets for Ma-T4 and Mc-T4 patients.

### Subgroups defined by fibroblast transcriptomes direct immunosuppressive phenotypes

We next analyzed fibroblast clusters of 69 ESCC datasets based on their transcriptomic similarity. The fibroblast clusters showed highly heterogenic features by individuals, which was not evident in T cells and myeloid cells (Fig. 5A). The correlation matrix of fibroblast identified five subgroups (F1, F2, F3, F4, and F5) of patients (Fig. 5B and 5C). Interestingly, most patients of the F4 subgroup overlapped with those of the T2 subgroup (Fig. 5D-5F). Meanwhile, the T4 subgroup characterized by abundant T_ex_ cells was mainly distributed to F1, F2, and F5 subgroups (Fig. 5D-5F). Therefore, we constructed combined F-T groups connecting fibroblast groups and T cell groups and compared them on UMAP, which showed that the F4-T2 was the most distinct subgroup on the UMAP of T cell (Fig. 5G). Furthermore, the F1-T4 subgroup expressed the highest level of T_ex_ cell markers compared to the others (Fig. 5H and Supplementary Fig. S4A). On the other hand, the F4-T2 group showed the most negligible expression of T_ex_ cell markers (Fig. 5H and Supplementary Fig. S4A). Based on these findings, we comparatively analyzed F1-T4 and F4-T2 sub-categorized fibroblasts using GSEA. Two hundred eleven signaling pathways overlapped in the F1-T4-positively significant dataset and the F4-T2-negatively significant dataset. Three signaling pathways coincided in the F1-T4-negatively significant dataset and F4-T2-positively significant dataset (Fig. 5I). Interestingly, among those overlapped signaling, we found interleukins and TGF-β signaling pathways were enriched in F1-T4 fibroblasts (Fig. 5I-5K). In βcontrast, complement process triggering signaling and FCGR (Fc-gamma receptor) activation signaling were enriched in F4-T2 fibroblasts. Additionally, F1-T4 and F4-T2 grouped T cells were analyzed by GSEA. Consistent with M-T classification, the F1-T4 group showed ‘PD-1 signaling’ and ‘IFN α/β signaling’ with positive NES while negative NES for F4-T2 T cells (Supplementary Fig. S4B). In epithelial cell analysis, ‘Mitochondrial electron transport NADH to Ubiquinone’ signaling was identified as specific to F1-T4 and F4-T2 with positive NES and negative NES, respectively, consistent with the results from M-T class analysis (Supplementary Fig. S4C). Notably, in comparison of M-T and F-T classification, we found all patients classified into Mb-T2 were also included in the F4-T2 group (Supplementary Fig. S4D). Although Ma-T4 and Mc-T4 classified patients were distributed to several groups of F-T class, F1-T4-, F4-T4-, and F5-T4-grouped patients only overlapped with Ma-T4 and Mc-T4 groups (Supplementary Fig. S4D).

**Figure 5.**
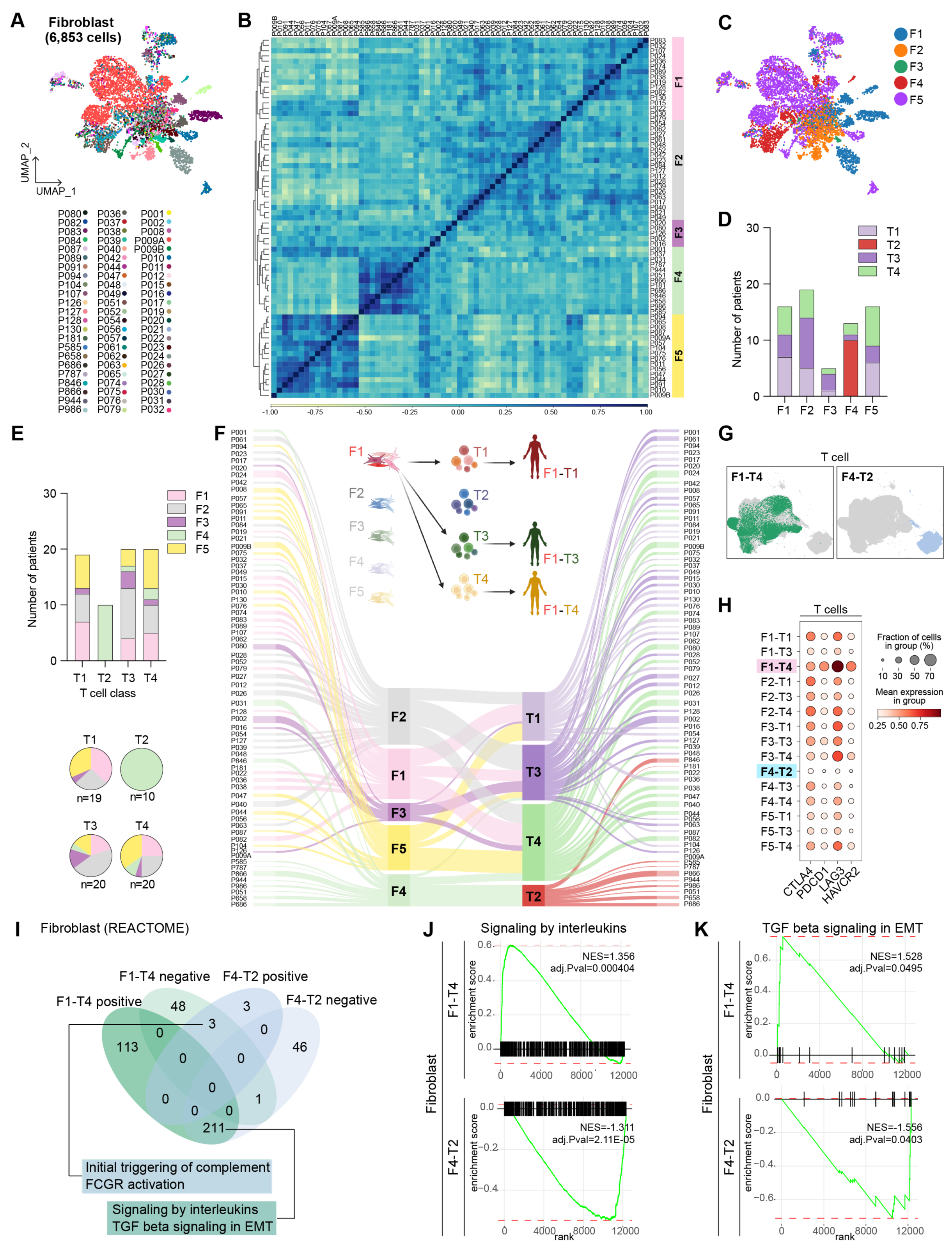
| Fibroblasts classification and patients grouping with T cell class. **A,** Fibroblasts of each patient were isolated and independently analyzed. UMAP labeled with each patient was shown. **B,** Fibroblasts were classified into five sub-groups by principal component analysis (PCA) and Pearson correlation. PCA result was clustered by the dendrogram, and Pearson correlation was displayed by color spectrum. **C,** Fibroblasts classification was displayed in UMAP. **D,** The number of patients of T cell sub-groups was displayed in each sub-group categorized by fibroblasts. **E,** The number of patients of fibroblast sub-groups was displayed in each sub-group categorized by T cells. The patient proportion of each fibroblast sub-group was shown on T cell sub-groups with pie plots. **F,** Sankey plot showing the connection of each patient’s fibroblast and T cell categories. Patients were re-grouped by fibroblast and T cell categories (F-T group), and T cells of the patients were shown with the F-T group. **G,** T_ex_ markers expression was compared in F-T groups in T cells with dot plots. **H,** Spatial location of T cells of F1-T4 and F4-T2 groups were shown on the UMAP. **I,** Fibroblasts from F1-T4 and F4-T2 groups were subjected to GSEA analysis using the REACTOME database. Significant signaling pathways were listed with positive values of NES and negative values of NES. Shared or exclusive signaling pathways between F1-T4 and F4-T2 were visualized with a Venn diagram. **J-K,** Overlapped signaling pathways in F1-T4-positive and F4-T2-negative values of NES from GSEA. Enrichment plots of Signaling by interleukins (**J**) and TGF-β signaling in EMT (**K**) were displayed.

In the further comparative cell-to-cell interaction analysis of F1-T4 and F4-T2- subgrouped patients, we found that overall, the number of signaling interactions between fibroblast and the other cell types was decreased in the F1-T4 subgroup compared to that of F4-T2 (Supplementary Fig. 4E). Collagen and integrin-mediated cell-to-cell interactions were primarily lost in the F1-T4 compared to F4-T2, which implicates plausible roles of collagen and integrin for immune cell activation by fibroblast (Supplementary Fig. S4F). We also analyzed the mast cells for the classification (Supplementary Fig. S5A). 3,993 cells were segregated from TME datasets and processed for calculating transcriptomic similarity (Supplementary Fig. S5B). However, the difference in transcriptomes represented by Pearson’s correlation was insufficient to make subgroups. Collectively, fibroblast and T cell-based classification identified the most significant differences in F1-T4 and F4-T2 groups with T_ex_ markers expression, IFN signaling in T cells, and TGF-β signaling in fibroblasts.

## Biomarkers of ESCC subtypes defined by TME transcriptomes

Our analyses found that patients of Ma-T4, Mc-T4, and F1-T4 subgroups have an immunosuppressive tumor niche, while patients of Mb-T2 and F4-T2 subgroups carry a tumor-unfavorable niche. Since the tumor niche is generated by the continuous interaction between tumor cells and TME, it is presumable that tumor niche-based classification also determines the characteristics of tumor cells. To determine distinct features of tumors in each group, we sought to identify markers for each group of patients and assess prognostic effects. In the epithelial cells, we first applied M (myeloid cells)-T (T cells) classification and found specific genes for Ma-T4 and Mc-T4 (ISG15, CFL1, PFN1, MYL6, MDK, and UBE2L6), and Mb-T2 (ASNSD1, RHOF, MRPL23, SNRPD3, and EIF3J) groups (Fig. 6A and 6B). Using F (fibroblasts)-T (T cells) classification, we also found F1-T4 (TPM2, MYL6, GABARAP, MRPL41, NDUFS8, and UBE2L6) and F4-T2 (SNRPD3, DNAJB9, EIF3J, EEF1G, and EIF1) specific genes (Fig. 6C and 6D). Intriguingly, we found immunosuppressive Ma-T4 and F1-T4 groups shared *MYL6* (Myosin Light Chain 6) and *UBE2L6* (Ubiquitin Conjugating Enzyme E2 L6) genes as biomarkers for tumor cells. Simultaneously, *SNRPD3* (Small Nuclear Ribonucleoprotein D3 Polypeptide) and *EIF3J* (Eukaryotic Translation Initiation Factor 3 Subunit J) genes were the overlapped biomarkers in tumor-favorable Mb-T2 and F4-T2 groups of epithelial cells (Fig. 6E-6G and Supplementary Fig. S6A-S6D).

**Figure 6.**
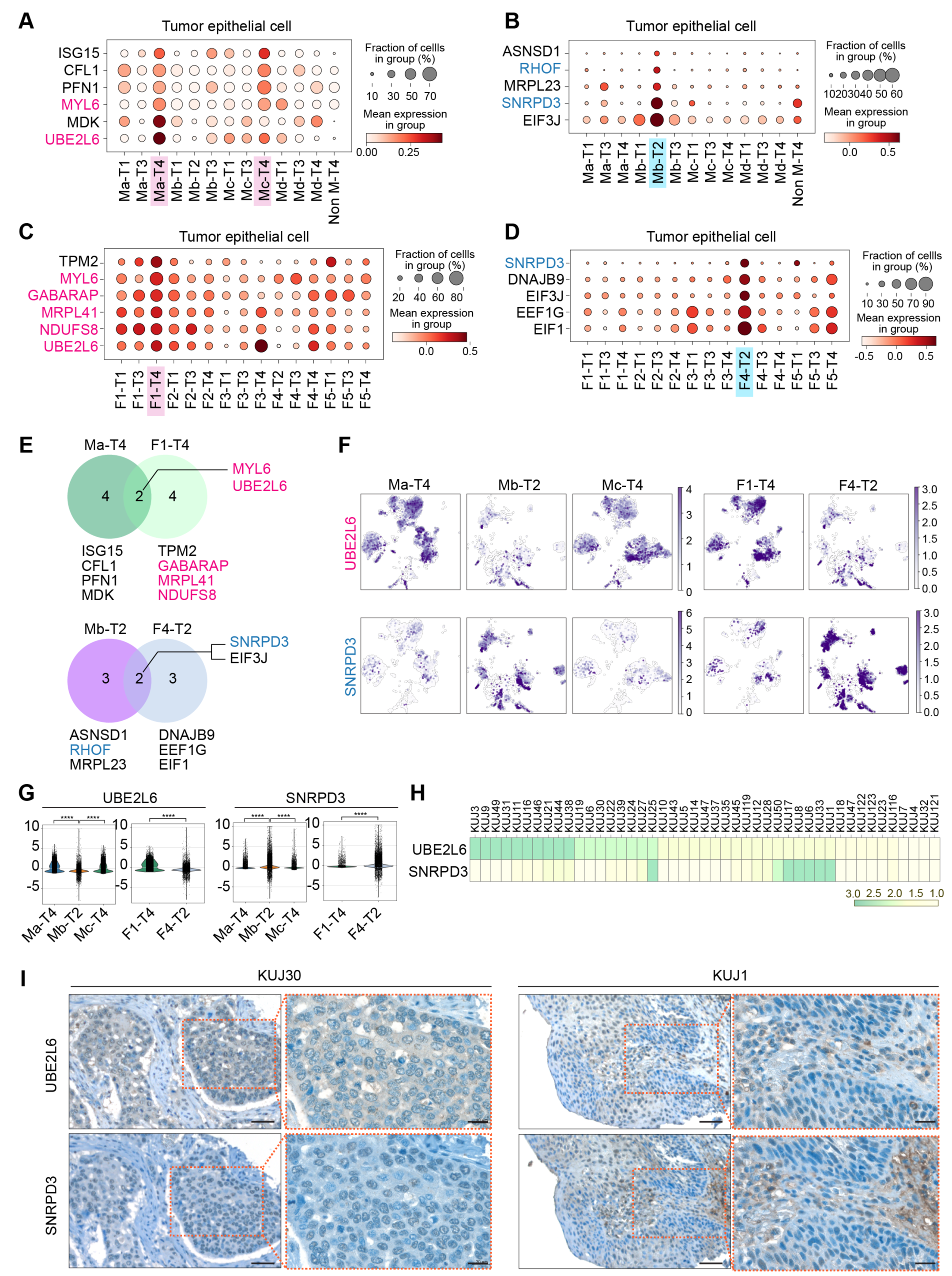
| Biomarkers of tumor cells based on M-T or F-T groups and their correlation with prognosis. **A-D,** Patients’ epithelial cells were grouped by M-T and F-T categories, and each group was projected to DEG analysis. The genes of which high expression are related to poor prognosis of ESCC patients were highlighted in red. The genes of which high expression related to better prognosis of ESCC patients were highlighted in blue. M1-T4- and M3-T4-specific (**A**) and M2-T2-specific (**B**) marker genes were displayed with dot plots. F1-T4-specific (**C**) and F4-T2-specific (**D**) marker genes were displayed with dot plots. **E-G,** Identified biomarkers from Ma-T4, F1-T4, Mb- T2, and F4-T2 were displayed with Venn diagram (**E**), and gene expression in each group was shown with UMAP (**F**). Expression of overlapped marker genes shown in the Venn diagram was compared in Ma-T4, Mb-T2, Mc-T4, F1-T4, and F4-T2 classified epithelial cells using violin plots (**G**). **H**-**I,** Immunohistochemistry of UBE2L6 and SNRPD3 from human ESCC were shown with scored heatmap (**H**) and representative images (**I**). IHC scores displayed from 1 (lowest expression) to 3 (highest expression). Scale bars = 50 μm (lower magnification) and 20 μm (higher magnification).****p<0.0001.

Then we determined the prognostic relevance of those specific genes using the TCGA database. Among the identified epithelial cell marker genes, we found that higher expression of *UBE2L6* and *MYL6* genes from Ma-T4 and Mc-T4 groups and *UBE2L6*, *MYL6*, *MRPL41*, *NDUFS8* (NADH:Ubiquinone Oxidoreductase Core Subunit S8), and *GABARAP* (GABA Type A Receptor-Associated Protein) genes from F1-T4 group was correlated with poor prognosis (Supplementary Fig. S6E). Meanwhile, ESCC patients with higher expression *RHOF* (Ras Homolog. Family Member F) and *SNRPD3* genes, markers of Mb-T2 or F4-T2, showed better prognosis (Fig. 6E and Supplementary Fig. S6E).

In addition to the biomarkers mainly expressed in tumor epithelial cells, we also tried to find markers expressed in myeloid cells, T cells, and fibroblasts from assigned subgroups. From the Ma-T4 and Mb-T2 subgroups in myeloid cells, *MS4A6A* (Membrane Spanning 4-Domains A6A) and *SNRPD3* were specifically expressed, respectively, with significant correlation with prognosis. Moreover, *SNRPD3* was repeatedly identified as an Mb-T2-subgrouped T cell biomarker (Supplementary Fig. S7A-S7E). Next, we identified markers specific to F1-T4 and F4-T2 subgroups of fibroblasts and T cells (Supplementary Figs. S7F-S7I). *S100A10* (S100 Calcium Binding Protein A10) and *FABP5* (Fatty Acid Binding Protein 5), F1-T4 grouped fibroblast specific markers, were correlated with poor prognosis (Supplementary Fig. S7J). In contrast, high expression of STK4 (Serine/Threonine Kinase 4), an F4-T2-grouped fibroblast-specific marker, was linked to a better prognosis (Supplementary Fig. S7J). Among the F1-T4 and F4-T2 subgroups-specific genes in T cells, high expression of *BAG3* (Bag Cochaperone 3) and *SNRPD3* were correlated with poor prognosis and better prognosis, respectively (Supplementary Fig. S7K). Interestingly, SNRPD3 was observed as a better prognostic marker in epithelial cells, myeloid cells, and T cells of the Mb-T2 subgroup, as well as epithelial cells and T cells of the F4-T2 subgroup. Therefore, it is likely that *SNRPD3* is a robust biomarker for patients with a tumor-unfavorable tumor niche. On the other hand, *UBE2L6* is expected to be a potent biomarker for patients with an immunosuppressive tumor niche, as higher expression of this gene was observed in the epithelial cells of the Ma-T4, Mc-T4, and F1-T4 subgroups. Based on these findings, we analyzed ESCC tumor microarray (TMA) samples to assess the expression of UBE2L6 and SNRPD3. UBE2L6 was highly expressed in 22.2 % (IHC score=3, n=10) of tumor samples, and SNRPD3 was markedly expressed in 13.3 % (IHC score=3, n=6) of patients (Fig. 6H and 6I). All UBE2L6^high^ patients showed a relatively lower expression of SNRPD3 (IHC score≤2), and 5 out of 6 SNRPD3^high^ patients displayed a low expression of UBE2L6 (IHC score≤2). As identified from datasets, UBE2L6 was detected mainly from tumor cells, while SNRPD3 staining was positive from TME and tumor cells. These results suggest that UBE2L6 and SNRPD3 are biomarkers exclusively expressed in ESCC patients, related to specific patient groups of immunosuppressive or tumor-unfavorable niches, respectively.

### Pathological relevance of TME transcriptomics to anti-PD-1 immunotherapy response

Since we have identified patient subgroups to predict the response to immunotherapy, we tested if our classification matches the immune cells of patients treated with anti-PD-1 immunotherapy. Patients were grouped into responders (R) and non-responders (NR) by their sensitivity to the PD-1 antibody treatment. Peripheral blood immune cells from three responders and three non-responders were collected to compare their phenotypes to our established classification. Cell types, including T cell, B cell, monocyte, neutrophil, and platelet, were annotated after integrating six datasets (Figs. 7A-7C). As expected, T_ex_ markers (TIGIT, HAVCR2, LAG3, and CTLA4) were observed to be highly expressed in responders than in non-responders, indicating that patients who are susceptible to ICB exhibit the higher T_ex_ signature in their PBMCs compared to the PBMCs of other patients (Fig. 7D and Supplementary Fig. S8A). Furthermore, TIGIT was significantly expressed in CD8 T cells of PBMCs in responders compared to non-responders (Figs. 7D-7F), consistent with our findings from Ma-T4 and Mc-T4 grouped patients. Therefore, we performed GSEA analysis in responders and non-responders from their T cell clusters. Then we compared the results with those from T cells of Ma-T4, Mc-T4, and F1-T4 groups. Interestingly, we found that ‘IFN signaling’ was significantly enriched in responders and Ma-T4, Mc-T4, and F1-T4 (Figs. 7G-7J). PD-1 signaling pathway-related genes were commonly enriched in Ma-T4 and F1-T4 groups of T cells. Moreover, T cell scoring analysis using PD-1 pathway genes and IFN pathway genes showed higher scores in responders, Ma-T4, Mc-T4, and F1-T4 groups compared to non-responders and Mb-T2 groups, respectively (Fig. 7K). These results echo the importance of IFN signaling in immunotherapy-sensitive patients, as we found from Ma-T4- and Mc-T4-grouped patients (Figs. 3G and 3I). ‘PD-1 signaling’ and ‘MHC class II antigen presentation’ of responders also overlapped with the GSEA results in Ma-T4/F1-T4 and Mc-T4 groups, respectively (Figs. 7G-7I and Supplementary Fig. S4B). Then we integrated single-cell transcriptomes of responders and non-responders with the transcriptomes of previously classified 13 M-T groups of 69 ESCC patients to test their transcriptomic proximity (Fig. 7L). From principal component analysis and Pearson’s correlation, the non-responder group was hierarchically closer to Mb-T2 than to Ma-T4 or Mc-T4. On the other hand, the transcriptome of responders showed a proximal cluster with Ma-T4 and Mc-T4 compared to Mb-T2 (Fig. 7M). A positive correlation between responders and Mc-T4 was evident when we narrowed down the comparison counterparts from 12 categories to 3 (Ma-T4, Mb-T2, and Mc-T4) categories (Supplementary Fig. 8B).

**Figure 7.**
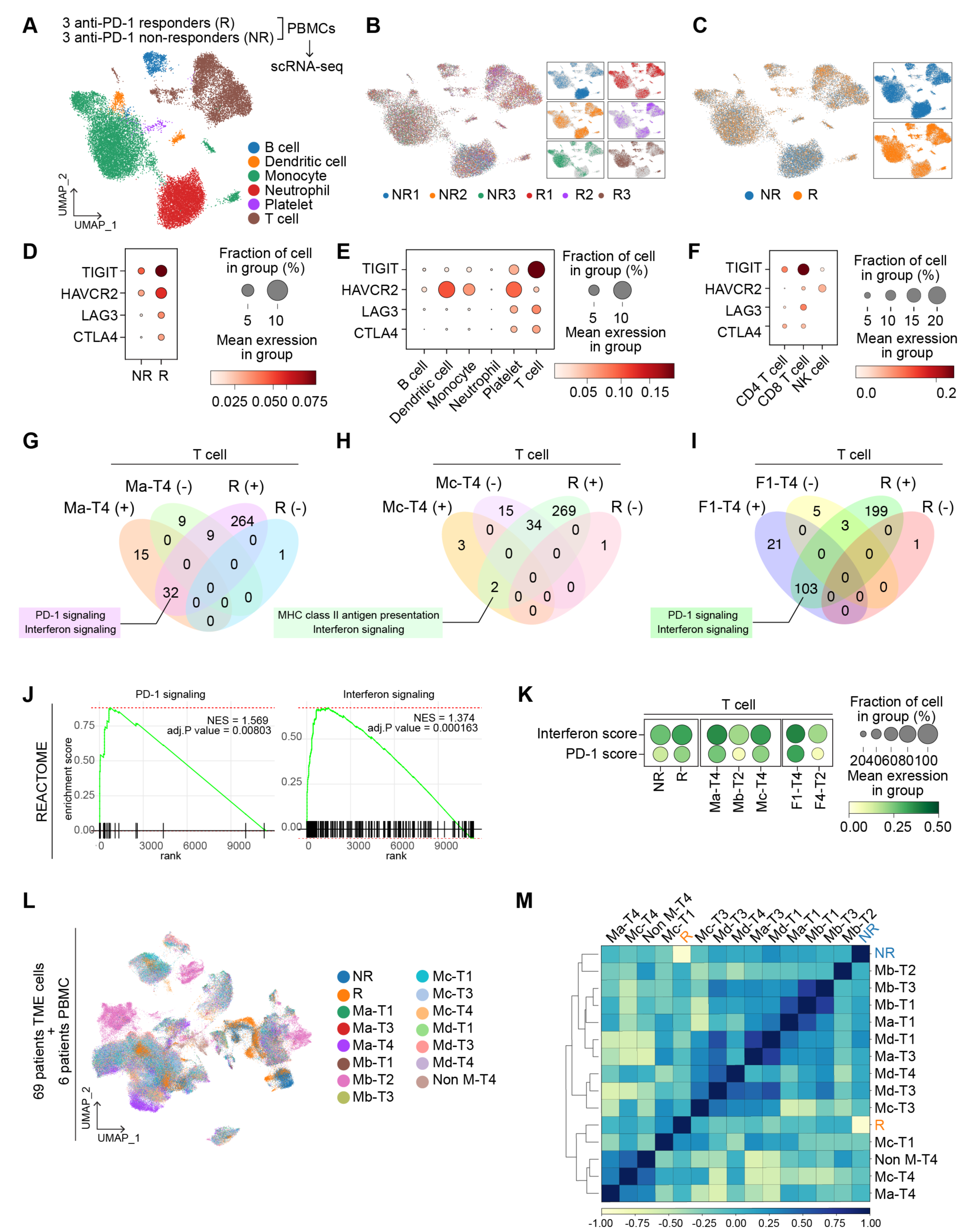
| Single-cell transcriptomics of immune cells of anti-PD-1 immunotherapy-treated patients. **A-C,** Peripheral blood immune cells transcriptomes of three responders (R) and three non-responders (NR) (to anti-PD-1 ICI) were integrated and presented with UMAP by cell types (**A**), patients (**B**), and response groups (R vs. NR) (**C**). **D-F,** T_ex_ marker genes expression were compared by the anti-PD-1 response (R vs. NR) (**D**), cell types (**E**), and T cell subsets (**F**). **G-J,** GSEA analysis performed by responders vs. non-responders using the REACTOME database. Significant results with positive NES and negative NES were listed with R positive and R negative, respectively. GSEA results were compared with Ma-T4 (**G**), Mc-T4 (**H**), and F1-T4 (**I**). Enrichment plots of PD-1 signaling and Interferon signaling were displayed (**J**). **K,** Pathway scores were compared in 3 groups (1) responders and non-responders, 2) Ma-T4, Mb-T2, and Mc-T4, 3) F1-T4 and F4-T2) and shown with dotplots. **L,** single-cell transcriptomes of immunotherapy-experienced patients were integrated with 69 ESCC patients’ TME transcriptomes and shown with UMAP by M-T groups and anti-PD-1 response groups. **M,** Correlation matrix with M-T patient groups and anti-PD-1 response groups. PCA result was clustered by the dendrogram, and Pearson correlation was displayed by color spectrum.

To evaluate the accuracy of our M-T classifications for immunotherapy, we compared the prediction results of different patient classifications with those of the M-T groups. Instead of the separated analyses, such as T cell only or myeloid cell only, post-integration subgrouping of T cell and myeloid cell clusters was performed to construct a new classification of patients (Supplementary Figs. S8C-S8D). After making this Myeloid and T cell-combined groups (MT1-MT11), we integrated the patients’ TME cells datasets with PBMCs datasets of responders and non-responders (Supplementary Fig. S8E). The new groups were not exclusive to the M-T groups, and the T_ex_ scores were the highest in the MT2 group in this new classification, while the scores were highest in the Ma-T4 and Mc-T4 in the M-T classification (Supplementary Figs. S8F-S8H). However, these new groups did not segregate responders and non-responders groups in the correlation analysis, indicating that none of the MT-combined groups (MT1 – MT11) showed a higher correlation with responders or non-responders groups than the proximity between responders and non-responders (Supplementary Fig. S8I). We next compared M-T classifications with patients grouped by T_ex_ cell markers expression. Using the T_ex_ cell scores in T cells, we grouped patients into four quartiles (High, High-Mid, Mid-Low, and Low). We integrated these data with responders and non-responders datasets (Supplementary Figs. S8J-S8L). Surprisingly, although most Ma-T4 and Mc-T4 patients were included in High or High-Mid, these T_ex_ cell markers-based groups did not show a positive correlation with responders (Supplementary Fig. S8M). The same workflow was used to classify patients based on the mean value (High and Low). However, the High group still did not show a positive correlation with responders (Supplementary Figs. S8N-S8Q). These results suggest that our M-T classifications are more accurate in predicting responders than myeloid-T cells-combined or T_ex_ cell markers-based categories.

## Discussion

To enhance the efficacy and minimize adverse effects of cancer therapies, it is crucial to subtype and characterize patients, selecting those who will benefit most from specific treatments. In this study, we curated a significant number of single-cell transcriptome datasets from human ESCC patients, establishing precise patient categories based on TME transcriptomes beyond conventional and molecular pathology (Fig. 8). We discovered that combining the transcriptional signatures of myeloid cells with T cells (M-T) or fibroblasts with T cells (F-T) can effectively stratify ESCC patients, predicting the outcomes of immunotherapy treatment. Specifically, patients classified as Ma-T4, Mc-T4, and F1-T4 displayed the T_ex_ cells phenotype in their T cells, suggesting a promising response to ICB. Conversely, patients categorized as Mb-T2 and F4-T2 were unlikely to respond to ICB, as their T cells rarely exhibited T cell exhaustion. The prediction of ICB efficacy was supported by comparing the transcriptomes of patients who had undergone immunotherapy, where Ma-T4, Mc-T4, F1-T4, and ICB-responders shared the same signature of IFN signaling, with Mc-T4 exhibiting close transcriptomic proximity to ICB-responders. Although current immunotherapy primarily focuses on T_ex_ cell markers, our M-T classification was expected to provide a better prediction for ICB response than grouping patients solely based on these markers.

**Figure 8.**
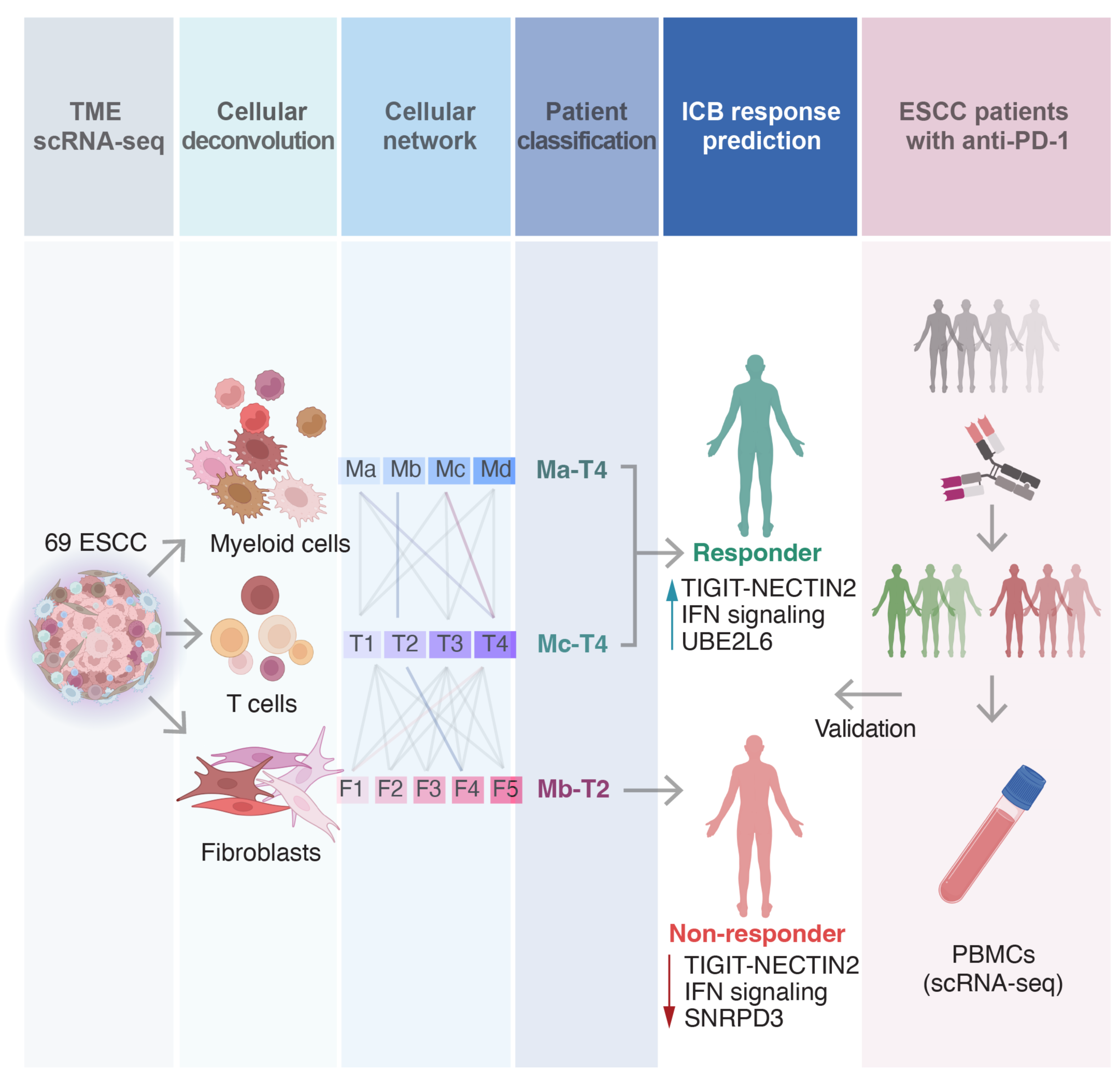
| Schematic representation of this study. Single-cell transcriptomes of ESCC patients’ TME cells were analyzed to predict immunotherapy response and identify biomarkers and potential adjuvant therapies to improve efficacy. The prediction of responsiveness was retrospectively validated by examining transcriptomes of ICB-experienced patients’ immune cells.

In addition to selecting patient groups for ICB response, we propose potential adjuvant therapies to improve ICB treatment efficacy. We have identified the NECTIN2-TIGIT axis as a significant interaction between tumor cells and immune cells in Ma-T4- or Mc-T4-grouped patients. NECTIN2 interacts with CD226 and TIGIT on the surface of T cells, with the latter acting as a competitive inhibitor of CD226.^33^ TIGIT prevents CD226 homodimerization by binding to CD226, which suppresses CD226-mediated T cell activation. ^34^ Moreover, TIGIT induces immunoregulatory effects by promoting the maturation of immunoregulatory dendritic cells, T_reg_ cells, and T_ex_ cells. ^35-38^ TIGIT in T_reg_ upregulates coinhibitory receptors such as HAVCR2/TIM-3, thus playing a critical role in immune responses. ^35^ Recent clinical trials have shown that TIGIT is a promising new target for ICB, with phase III clinical trials for esophageal cancer (skyscraper-07 and skyscraper-08) ongoing. However, phase II and III clinical trials with lung cancer patients have generated mixed results, ^39, 40^ with the latest results showing improved overall response rate (ORR) and progression-free survival (PFS) when the anti-TIGIT antibody was combined with the anti-PD-1 antibody in NSCLC patients, ^40^ but not in patients with extensive-stage small-cell lung cancer (ES-SCLC). ^39^ Our findings suggest that patient selection for ICB treatment needs to be based on additional standards beyond PD-1 or PD-L1/2 expression in tumors. Categorizing patients into detailed groups based on TME transcriptomes may improve the efficacy of ICB treatment.

To enhance immunotherapy’s efficacy, targeting enriched signaling pathways in each group would also be promising. For example, suppression of IFN signaling in Ma-T4- and Mc-T4-like patients and TGF-β signaling or interleukin signaling in F1-T4-like patients might improve ICB efficacy. Although a majority of studies focused on the roles of the anti-tumor effect of IFN α/β, a recent study revealed that type I IFN protects cancer cells from T cell-mediated cytotoxicity.^41, 42^ Furthermore, persistent IFN signaling activation induces resistance to ICB therapy.^41, 43^ Accordingly, ISG15, an IFN-stimulated gene, and UBE2L6 are highly expressed in the tumor cells of Ma-T4- or Mc-T4-grouped patients, as observed in other types of cancer.^44, 45^ Considering the roles of ISG15 and UBE2L6 in controlling TP53 stability by ISGylation, IFN signaling enriched groups might experience frequent intratumoral genetic alterations through the downregulation of TP53.^46, 47^ Notably, IFN signaling is specifically activated in the T cells of PD-1 immunotherapy responders as well as those of Ma-T4, Mc-T4, and F1-T4 (Fig. 7). These results highlight the robustness of IFN signaling across tumor cells and immune cells in immunotherapy-applicable patient groups, which can serve as an adjuvant target for immunotherapy.

While our results primarily rely on the transcriptional networks of the TME, we did not include the transcriptional signatures of tumor cells in identifying biomarkers. Nevertheless, we identified biomarkers in patient groups despite the inter-tumoral heterogeneity, suggesting that the expression of these biomarker genes may be associated with TME-released factors. Notably, IFN-stimulated genes such as ISG15 and UBE2L6 were among the identified biomarkers. In addition to in silico analyses, future studies are needed to determine the therapeutic impact of anti-TIGIT pathway inhibitors or IFN signaling inhibitors combined with ICBs on a specific group of esophageal squamous cell carcinoma patients. Furthermore, examining more datasets from ICB-experienced patients beyond the six reference datasets (responders and non-responders) we used here will provide a strong demonstration of the accuracy of our classification.

This study stratifies ESCC patients based on myeloid, T cell, and fibroblast transcriptome analysis and proposes potential adjuvant targets to improve cancer immunotherapy for specific subtypes of ESCC. Additionally, utilizing the most extensive reference of ESCC TME transcriptome, this study provides new insight into tumor niche remodeling in ESCC.

## Acknowledgments

We thank Pierre D. McCrea and Malgorzata Kloc for their insightful comments and the Herbert Irving Comprehensive Cancer Center for the shared resources (Biostatistics, Genomics, and Molecular Pathology). This work was supported by the Cancer Prevention and Research Institute of Texas (RP200315 to J.-I.P.), the National Cancer Institute (CA193297 and CA256207 to J.-I.P; 5P30CA013696 and 5P01CA098101 to A.-K.R., H.N., K.D., G.E., C.M.), an Institutional Research Grant (MD Anderson to J.- I.P.), a Specialized Program of Research Excellence (SPORE) grant in endometrial cancer (P50 CA83639), and Radiation Oncology Research Initiatives. Schematic representation was created with Biorender.com.

## Author contributions

K.-P.K. and J.-I.P. conceived and designed the experiments. K.-P.K., S.Z., Y.H., B.K., G.Z., S.J., and J.Z. performed the experiments. K.-P.K., H.N., H.Z., and J.-I.P. analyzed the data. C.M., K.J.D., G.E., A.-K.R., and H.N. provided the ESCC TMA slides. H.Z. provided the single-cell RNA-seq datasets of PD-1-treated patients. K.-P.K. and J.-I.P. wrote the manuscript.

## STAR Methods

### RESOURCE AVAILABILITY

#### Lead contact

Additional information and requests for resources and reagents should be directed to and will be fulfilled by the Lead Contact, Jae-Il Park.

#### Materials availability

The materials will be available upon request.

#### Data and code availability

scRNA-seq data are available via the National Center for Biotechnology Information Sequence Read Archive (SRA) under the accession numbers PRJNA777911 and PRJNA672851. The code used to reproduce the analyses described in this manuscript can be accessed via GitHub (https://github.com/jaeilparklab/ESCC_project_2) and is available upon request.

### METHOD DETAILS

#### scRNA-seq data preparation

##### Public datasets

The raw read files of ESCC patient datasets were downloaded using the parallel-fastq-dump package and converted to fastq files. The fastq files were mapped to the GRCh38 reference genome using CellRanger (v7.0.1) pipeline. The datasets from 9 patients (NCBI BioProject: PRJNA777911) were utilized to CellRanger directly, while 60 patients’ datasets (NCBI BioProject: PRJNA672851) were separately input to CellRanger as CD45+ and CD45-datasets were sorted during sample preparation. Single-cell dataset and patient information are described in Supplementary Table 1.

#### scRNA-seq data analysis

##### Integration and clustering

The datasets from 9 patients were preprocessed independently, and the CD45^+^ cell clusters were retained for the immune cell population. 60 patients’ dataset analysis was started with CD45^+^ sorted datasets. After preprocessing procedures, 11 patients and 58 patients datasets were integrated using the “concatenate” function in Scanpy. A batch correction was conducted using “Harmony” implemented in Scanpy.^24^ “Louvain” algorithm was used for clustering cells. Each cell cluster was annotated primarily with “B cell”, “Fibroblast”, “Mast cell”, “Myeloid cell”, and “T cell” using marker genes of each cluster. T cells were further annotated with “CD4 T cell”, “CD8 T cell”, “exhausted T cell”, and “effector T cell” and Myeloid cells were further annotated into “Monocyte”, “Macrophage”, “M1 Macrophage”, and “M2 Macrophage” clusters.

##### Classification of each cell type

“T cell”, “Myeloid cell”, “Fibroblast”, and “Mast cell” clusters were isolated, and each cell type was analyzed with individual patients. The transcriptomic similarity of each patient was compared using the correlation matrix function in Scanpy. Dendrograms were drawn to show PCA proximity, and Pearson correlation was displayed with color code. Patients were clustered and classified based on the result of the correlation matrix. Patients were classified by T cell, myeloid cell, and fibroblast transcriptomes-based categories, then connected classifications such as Myeloid cell-T cell (M-T) and Fibroblast-T cell (F-T) were applied to each patient. The connected classification of each patient was visualized with a Sankey plot using the “pysankey2” package.

##### Cell-to-cell interaction analysis

“CellChat” package was used for the cell-to-cell interaction inference. To acquire intercellular interactions, epithelial cell datasets were added to immune cell datasets. For 11 patient datasets, excluded CD45-cell clusters were re-integrated into the immune cell datasets. For 58 patients’ datasets, CD45^+^ datasets were analyzed from separated matrix files. After preprocessing epithelial cells, “epithelial cells”, “effector T cell”, “exhausted T cell”, “CD4 T cell”, “CD8 T cell”, “M1 macrophage”, “M2 macrophage”, “Macrophage”, “Monocyte”, “B cell”, “Fibroblast”, and “Mast cell” clusters were merged. M-T or F-T classification-based patient groups were used to generate gene expression matrices for the CellChat analysis. From significant signaling pathways, “TIGIT” and “NECTIN” signaling were specified for analysis in each group of patients. Comparative analysis was performed using two different groups of patients (Ma-T4 vs. Mb-T2 and Mc-T4 vs. Mb-T2).

##### fGSEA analysis

“fGSEA” package was used for the GSEA analysis of Ma-T4, Mb-T2, Mc-T4, F1-T4, and F4-T2 groups of patients. “Epithelial cells”, “Myeloid cell”, “T cell”, and “Fibroblast” clusters were independently analyzed to obtain a differentially expressed gene (DEG) list. DEG was performed in Scanpy with the “rank_gene_groups” function using the “Wilcoxon” method. “C2” category and “REACTOME” subcategory or “C5” category and “GO:BP” subcategory were used to use each database. GSEA results are listed in Supplementary Tables 2 - Supplementary Table 5.

#### PBMCs scRNA-seq data analysis

##### Integration and clustering

PBMCs scRNA-seq datasets from anti-PD-1 therapy responders and non-responders were provided by Dr. Haiyang Zhang.^48^ Three responders’ and three non-responders’ gene expression matrix files were independently preprocessed and integrated. The batch effect was reduced by Harmony algorithm^24^, and cell types were annotated with markers used in the previous study.^48^ PBMC datasets were further integrated with 69 patients’ human ESCC datasets with the same workflow and analyzed.

##### fGSEA analysis

fGSEA analyses were performed with isolated T cells with DEG lists between responders and non-responders, as described above. REACTOME database was used, and the results were compared with human ESCC patient fGSEA results. GSEA results of PBMCs are listed in Supplementary Table 6.

##### Pathway score analysis

The Pathway scores were performed using the “scanpy.tl.score_genes” function implemented in the Scanpy package. The analysis were done with default parameters and the reference genes from MSigDB and other literature. Reference genes were listed in Supplementary Table 7.

#### Kaplan-Meier analysis

The survival of ESCC patients was analyzed based on the gene expression using the publicly available database (Kaplan-Meier Plotter, http://kmplot.com/analysis). Survival data of 81 ESCC patients were classified into two groups by gene expression. Patients were labeled with “high” when the expression of a gene of interest was above the median expression value, and the others were labeled with “low”. The survival of patient groups was compared until 70 months, and hazard ratio (HR) and the log-rank *P* value (logrank P) were indicated.

#### Immunohistochemistry

Immunostaining was performed as previously described.^49^ ESCC cancer tissue microarray slides containing 144 samples from 55 patients were provided by Dr. Hiroshi Nakagawa. Antigens were retrieved from paraffin-embedded tissues using a basic (pH 9.0) buffer. After blocking the tissues in PBS with goat serum, samples were incubated with primary antibodies (UBE2L6 [1:200] and SNRPD3 [1:200]) Detection was performed using an HRP-conjugated secondary antibody, followed by DAB. Samples were counterstained with hematoxylin and mounted with coverslips. The immunohistochemistry results were scored from 0 to 3, then analyzed and visualized using R and GraphPad Prism (v9.2.0).

#### Statistical analysis

The Student’s *t*-test was used to compare two groups (n ≥ 3), and a one-way analysis of statistical variance evaluation was used to compare at least three groups (n ≥ 3). *P* values < 0.05 were considered significant. Error bars indicate the standard deviation (s.d.). All experiments were performed three or more times independently under identical or similar conditions.

## Supplementary Information

**Supplementary information titles**

**Supplementary Figure 1.** Subtypes of T cells and myeloid cells.

**Supplementary Figure 2.** Myeloid-T cell (M-T) classification-based analysis in different cell types.

**Supplementary Figure 3.** Epithelial cell analysis by myeloid-T cell (M-T) classification.

**Supplementary Figure 4.** Cell-to-cell interactions comparison in F1-T4 and F4-T2 groups.

**Supplementary Figure 5.** Transcriptomic analysis of mast cell cluster.

**Supplementary Figure 6.** M-T groups-and F-T groups-based biomarkers of TME cells and their correlation with prognosis.

**Supplementary Figure 7.** M-T groups-and F-T groups-based biomarkers of TME cells and their correlation with prognosis.

**Supplementary Figure 8.** Comparison between responders and non-responders for anti-PD-1 immunotherapy.

**Supplementary Table 1.** The information of single-cell RNA-seq datasets. Supplementary Table 2. GSEA results of T cells of ESCC patients by M-T classifications, related to Figure 3.

**Supplementary Table 3.** GSEA results of epithelial cells and myeloid cells of ESCC patients by M-T classifications, related to Figure S2.

**Supplementary Table 4.** GSEA results of fibroblasts of ESCC patients by F-T classifications, related to Figure 5.

**Supplementary Table 5.** GSEA results of T cells and epithelial cells of ESCC patients by F-T classifications, related to Figure S4.

**Supplementary Table 6.** GSEA results of T cells of anti-PD-1 responders and non-responders, related to Figure 7.

**Supplementary Table 7.** Gene lists for score analysis, related to Figure 7.

## Supplementary Figure Legends

**Supplementary Figure 1.**
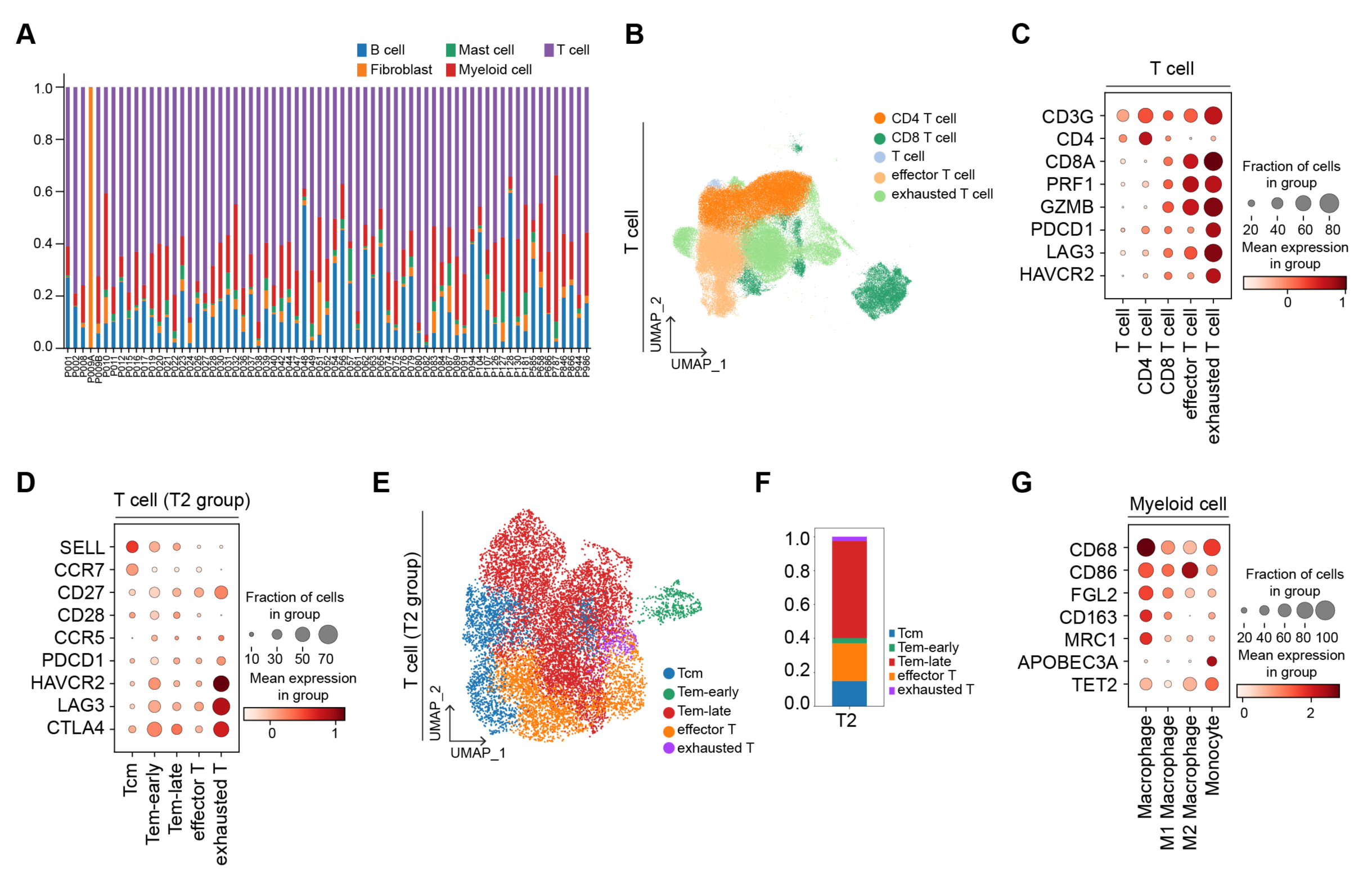
| Subtypes of T cells and myeloid cells. **A,** Proportion of cell types in each patient. **B,** UMAP of T cells with subset clusters. **C,** Marker genes of T cell subset expression were displayed with a dot plot. **D,** Detailed subsets of the T2 sub-group were shown with marker genes. **E-F,** Detailed subsets in the T2 sub-group were shI with UMAP (**E**) and stacked bar plot (**F**). **G,** Subsets of myeloid cells were shown with marker genes expression using a dot plot.

**Supplementary Figure 2.**
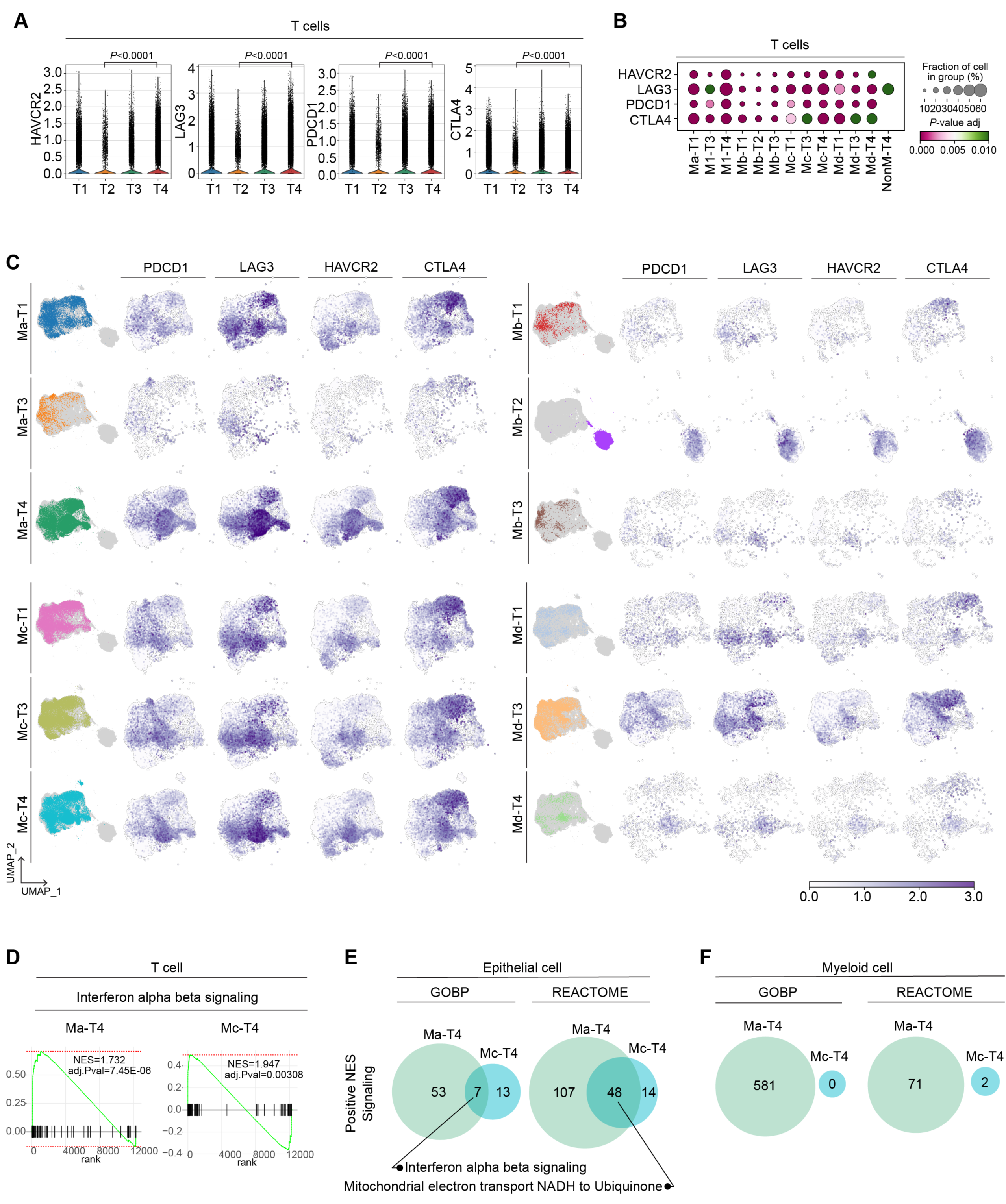
| Myeloid-T cell (M-T) classification-based analysis in different cell types. **A,** T_ex_ cell marker genes expressions in T cell-based sub-groups from T cell cluster. **B,** Dot plot showing T_ex_ cell marker genes expression and significance in M-T classified T cells. Each group was analyzed with the rest for each gene expression, and t-test results were displayed with color spectrum. **C,** T_ex_ cell markers expression in each M-T sub-group of T cells. **D,** Enrichment plot of ‘Interferon alpha beta signaling’ pathway from GSEA analysis of Ma-T4- and Mc-T4-grouped T cells. **E,** The results of GSEA analysis of epithelial cells in Ma-T4 and Mc-T4 groups of patients were compared. GOBP and REACTOME databases were used, and the significant signaling with a positive value of NES was compared. Overlapped signaling pathways were displayed with a Venn diagram. **F,** Myeloid cells of Ma-T4 and Mc-T4 groups were analyzed with GSEA using GOBP and REACTOME database. Significant signaling with a positive value of NES was listed and displayed with a Venn diagram.

**Supplementary Figure 3.**
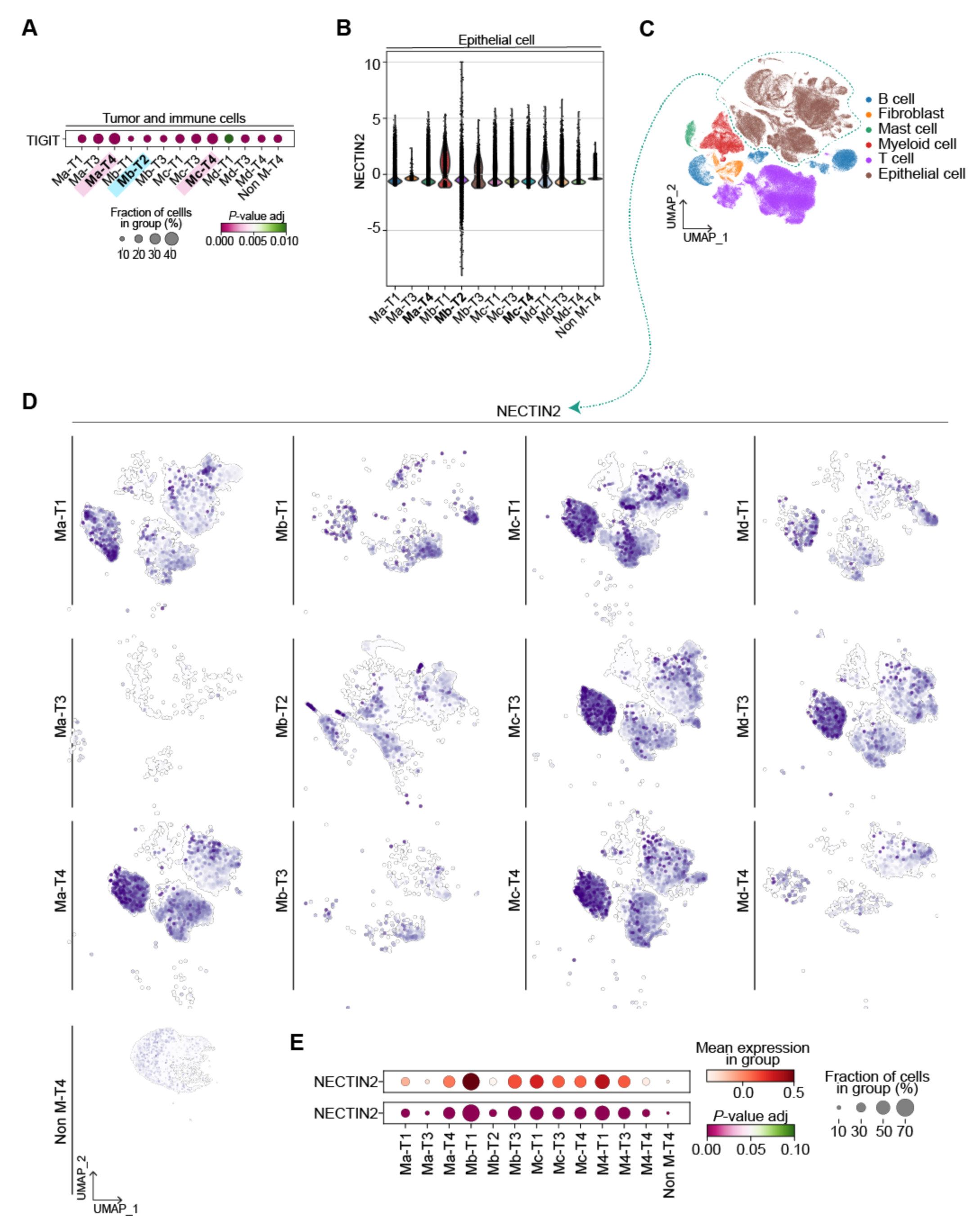
| Epithelial cell analysis by myeloid-T cell (M-T) classification. **A,** Dot plot showing the TIGIT expression in M-T classified epithelial cells and its significance. TIGIT expression in each group was compared to the rest of the groups to conduct a *t*-test. **B,** NECTIN2 expression in epithelial cells, grouped by M-T classification. **C,** UMAP display with whole cells. Epithelial cell cluster is highlighted. **D,** NECTIN2 expression comparison in M-T classified epithelial cells. **E,** NECTIN2 expression and significance in each M-T classified epithelial cell group. NECTIN2 expression in each group was compared to the rest of the groups to conduct a *t*-test.

**Supplementary Figure 4.**
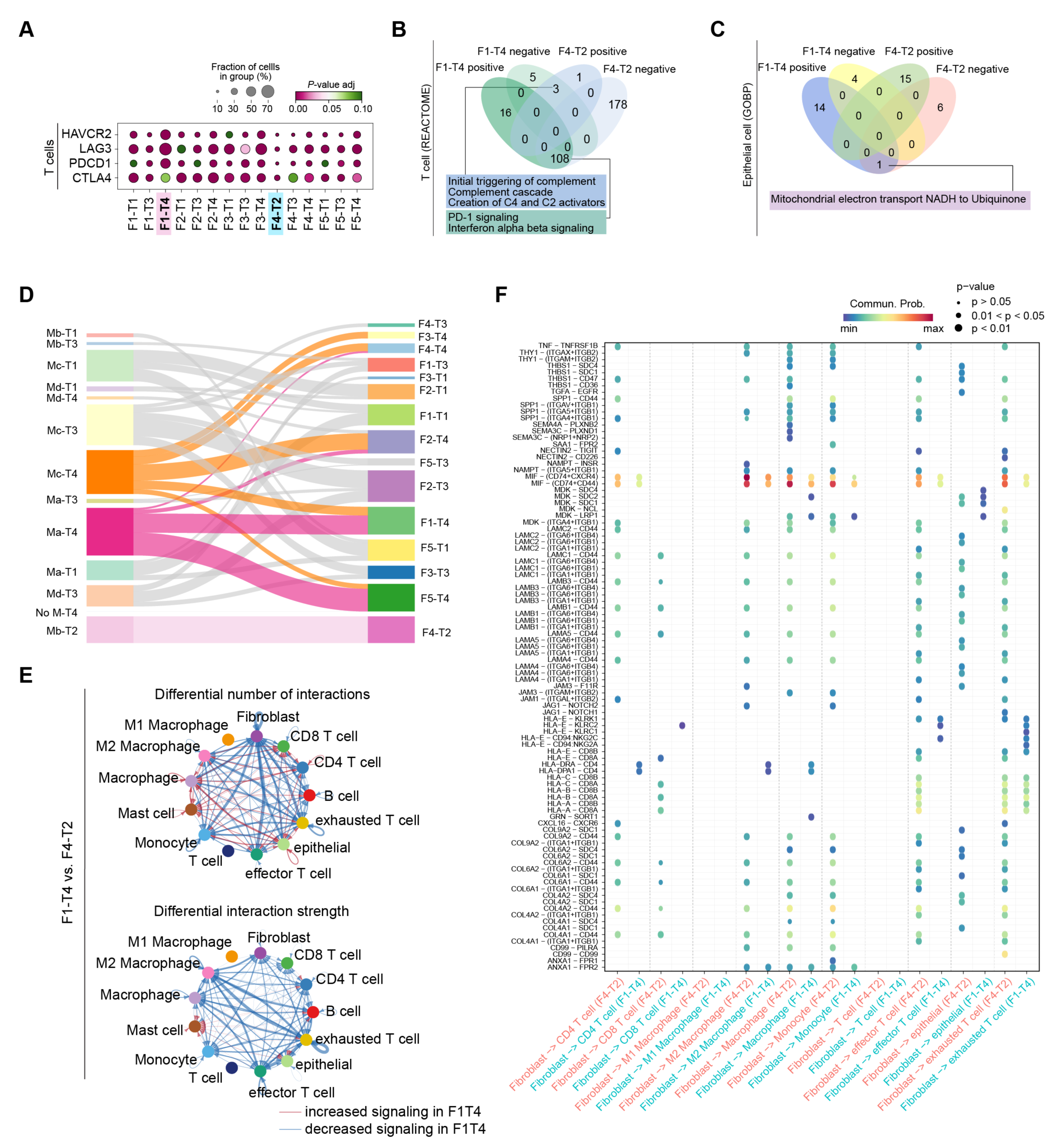
| Cell-to-cell interactions comparison in F1-T4 and F4-T2 groups. **A,** T_ex_ marker genes expression in fibroblast-T cell (F-T) classified T cells. The fraction of cells in the group expressing each gens and significance are displayed. Each gene in the sub-group was analyzed with the rest of the groups to perform a *t*-test. **B,** T cells grouped by F1-T4 and F4-T2 were subjected to GSEA using the REACTOME database. A list of pathways with positive or negative NES was displayed with a Venn diagram. **C,** GSEA results from epithelial cells of F1-T4 and F4-T2 were listed and compared with the Venn diagram. **D,** Patients of myeloid-T cells (M-T) and fibroblast-T cells (F-T) were compared using the Sankey plot. **E,** Comparative circle plot showing the significant signaling in F1-T4 and F4-T2 groups. The total number of interactions (top) and interaction weight (bottom) were compared in two groups. Red lines indicate increased signaling in F1-T4, and blue lines show decreased signaling in F4-T2. **F,** Interactions between fibroblast and other cell types were calculated and compared in F1T4 and F4-T2 groups. The sources (or ligands) from fibroblast and Receiver (or receptor) of different cells were displayed with a bubble plot.

**Supplementary Figure 5.**
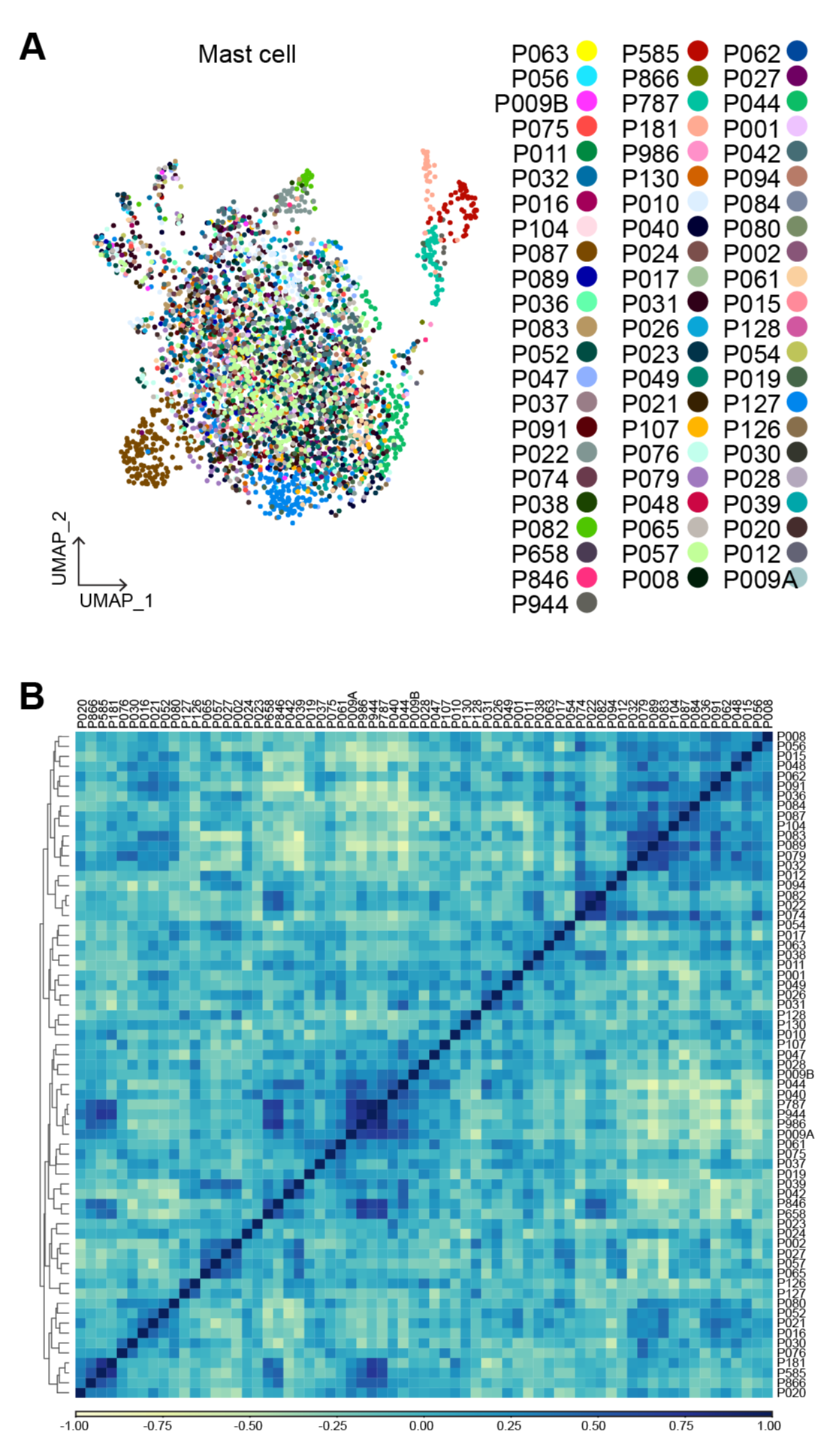
| Transcriptomic analysis of mast cell cluster. **A,** Mast cells were isolated from non-epithelial cells, and UMAP was re-drawn with individual patient information. **B,** Fibroblasts were analyzed by principal component analysis (PCA) and Pearson correlation. PCA result was clustered by the dendrogram, and Pearson correlation was displayed by color spectrum.

**Supplementary Figure 6.**
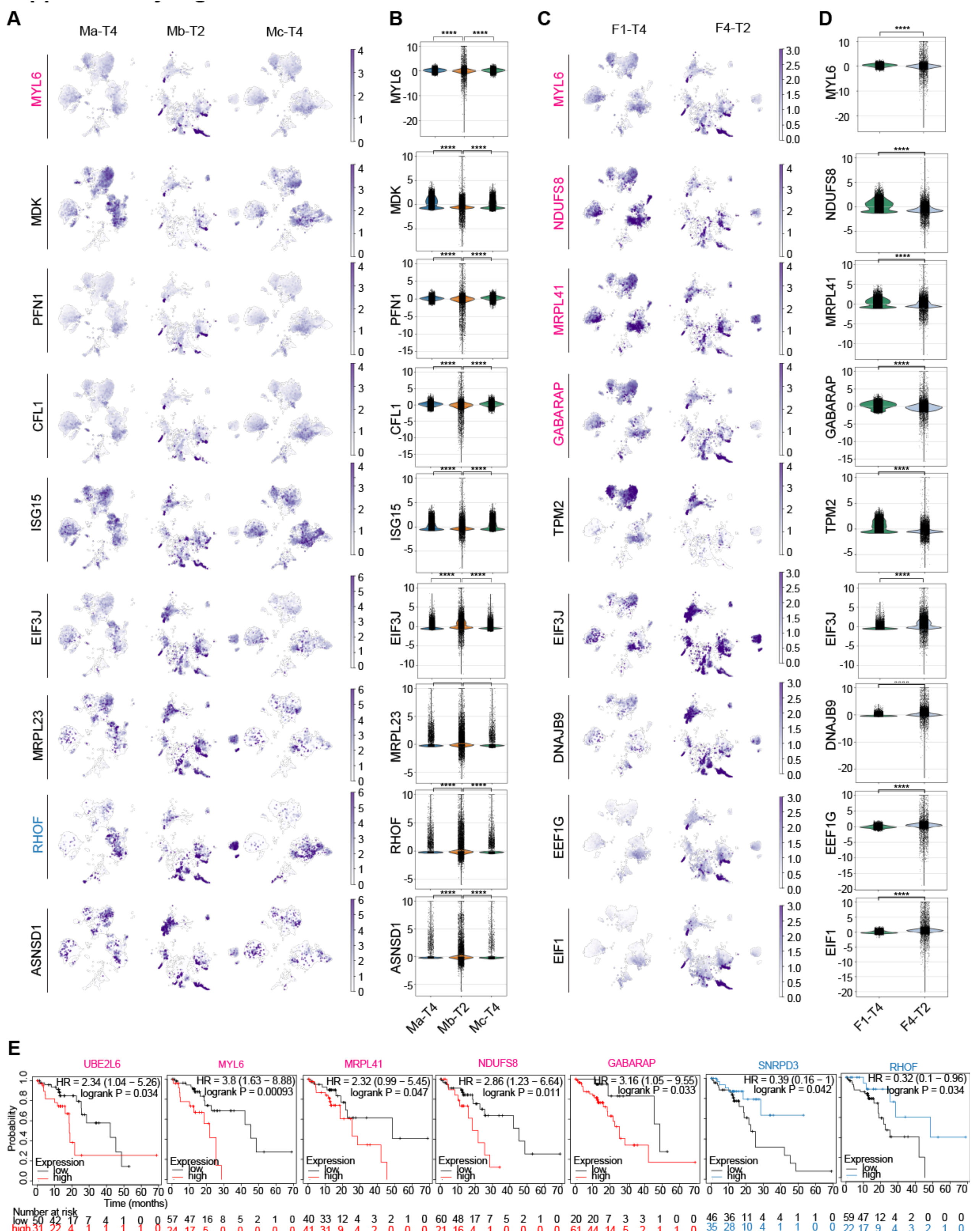
| M-T groups-and F-T groups-based biomarkers of TME cells and their correlation with prognosis. **A-B,** Expression of marker genes of epithelial cells in Ma-T4, Mc-T4, and Mb-T2 groups were shown with UMAP (**A**) and violin plots (**B**). **C-D,** Expression of markers from F1-T4 and F4-T2 classified epithelial cells with UMAP (**C**) and violin plots (**D**). **E,** Markers from epithelial cells from Ma-T4, Mb-T2, F1-T4, and F4-T2 categories were evaluated with their prognostic correlation of ESCC patients using Kaplan-Meier plots. HR, Hazard Ratio. ****p<0.0001.

**Supplementary Figure 7.**
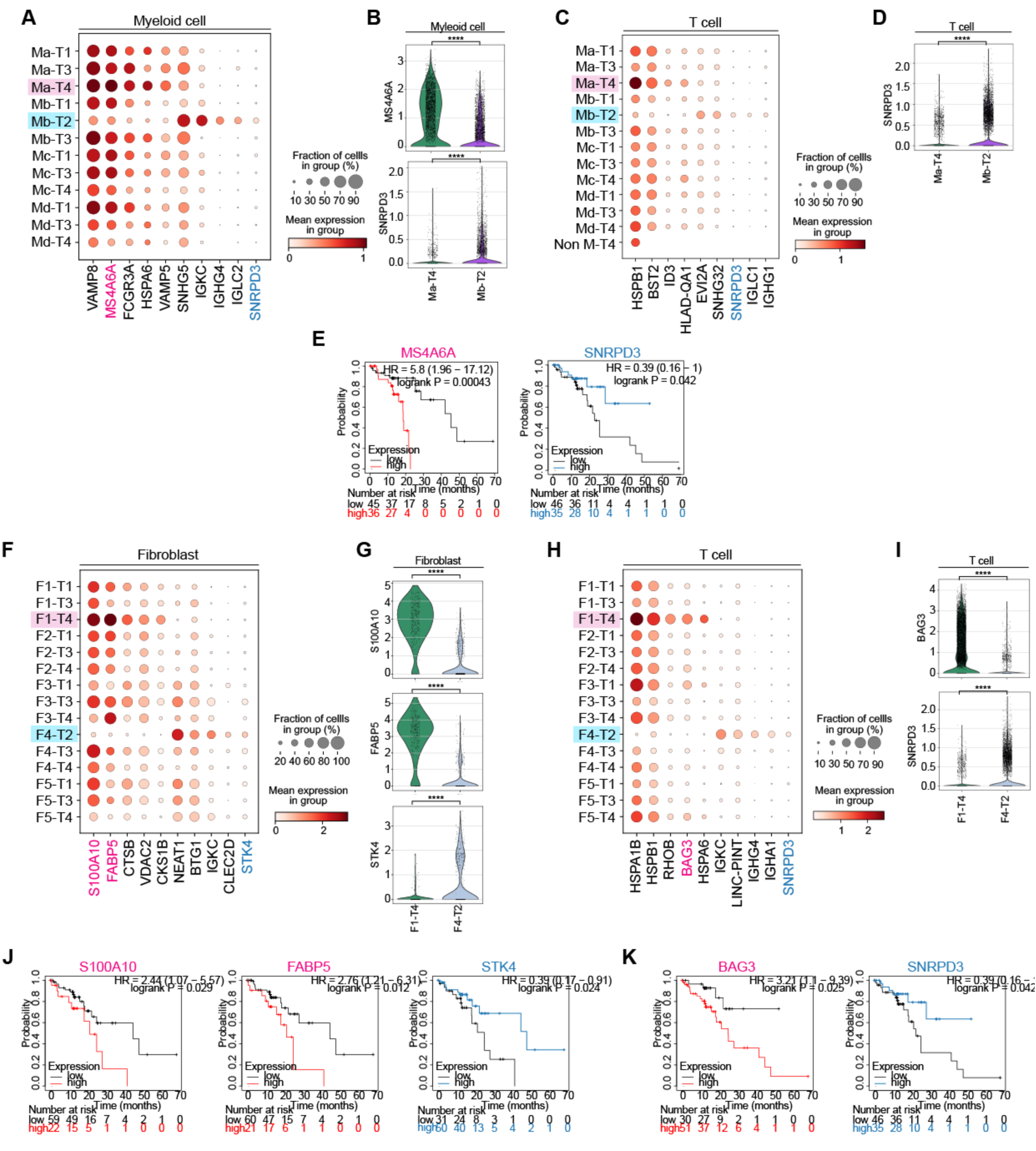
| M-T groups-and F-T groups-based biomarkers of TME cells and their correlation with prognosis. **A-B,** Markers of myeloid cells in Ma-T4 and Mb-T2 groups of patients were shown with dot plot (**A**), and prognostically correlated genes were shown with violin plot (**B**). **C-D,** Markers of T cells in Ma-T4 and Mb-T2 groups of patients were displayed with dot plot (**C**), and prognostically correlated gene was shown with violin plot (**D**). **E,** Ma-T4 and Mb-T2 groups-specific markers in myeloid and T cells were evaluated with their prognostic correlation of ESCC patients using Kaplan-Meier plots. **F-G,** Markers of fibroblasts in F1-T4 and F4-T2 groups of patients were shown with dot plot (**F**), and prognostically correlated genes were shown with violin plot (**G**). **H-I,** Markers of T cells in F1-T4 and F4-T2 groups of patients were displayed with dot plot (**H**), and prognostically correlated genes were shown with violin plot (**I**). **J,** Markers of F1-T4 and F4-T2 groups of fibroblasts were evaluated with their prognostic correlation of ESCC patients. Kaplan-Meier plots were displayed with an F1-T4 marker (S100A10 and FABP5) and an F4-T2 marker (STK4). **K,** Markers of F1-T4 and F4-T2 T cell groups were analyzed for their prognostic correlation in ESCC patients. F1-T4 marker (BAG3) and F4-T2 marker (SNRPD3) were displayed with Kaplan-Meier plots. HR, Hazard Ratio.

**Supplementary Figure 8.**
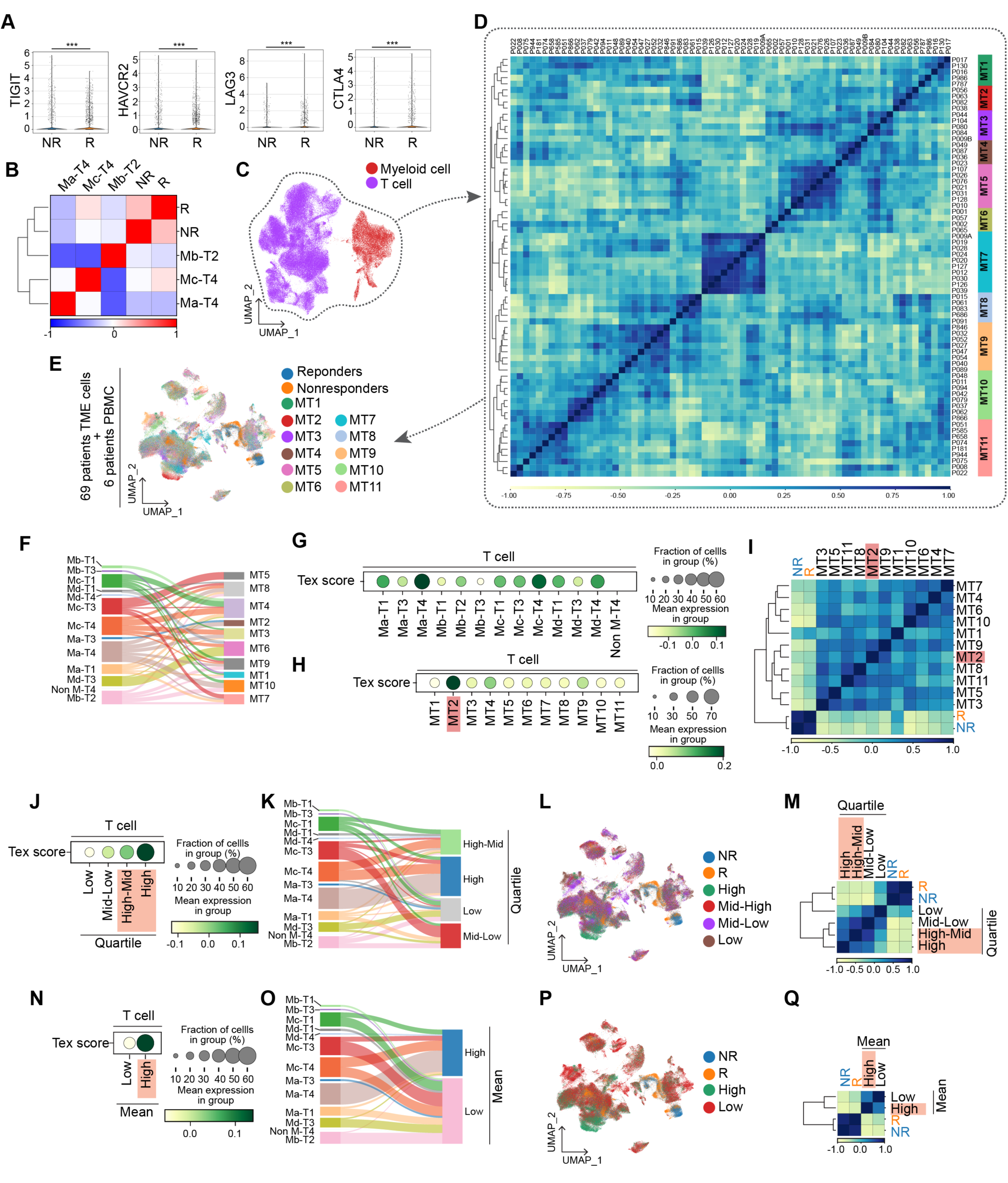
| Comparison between responders and non-responders for anti-PD-1 immunotherapy. **A,** T_ex_ markers expression was compared between responders and non-responders groups. ****p<0.0001. **B,** Correlation matrix with three M-T patient groups (Ma-T4, Mb-T2, and Mc-T4) and anti-PD-1 response groups (R and NR). PCA result was clustered by the dendrogram, and Pearson correlation was displayed by color spectrum. **C-D,** T cell and myeloid cell clusters of 69 ESCC patients were integrated first (**C**) and subjected to correlation analysis to find 11 myeloid-T cell-combined subgroups (**D**). **E,** TME transcriptomes of myeloid-T cell combined subgroups were integrated with PBMCs transcriptomes of anti-PD-1 responders and non-responders. **F,** Myeloid-T cell-combined subgroups were compared with M-T classifications using the Sankey plot. **G-H,** exhausted T cell scores were assessed in M-T groups (**G**) and myeloid-T cell-combined subgroups (**H**). **I,** Correlation matrix with myeloid-T cell-combined subgroups and anti-PD-1 response groups. PCA result was clustered by the dendrogram, and Pearson correlation was displayed by color spectrum. **J-K,** 69 ESCC patients were categorized into 4 quartiles by exhausted T cell score values (High, High-Mid, Mid-Low, and Low) in T cells (**J**) and compared with M-T groups (**K**). **L-M,** ESCC patients’ transcriptomes grouped by quartile values were integrated with transcriptomes of anti-PD-1 responders and non-responders (**L**), followed by correlation analysis (**M**). N-O, Transcriptomes of 69 ESCC patients were divided into 2 groups by mean value of exhausted T cell scores in T cells (**N**) and compared with M-T groups (**O**). **P-Q,** 69 ESCC patients’ TME transcriptomes were integrated with PBMCs transcriptomes from anti-PD-1-experienced patients (**P**), and subjected to correlation analysis (**Q**).

